# Toward single particle reconstruction without particle picking: Breaking the detection limit

**DOI:** 10.1101/431080

**Authors:** Tamir Bendory, Nicolas Boumal, William Leeb, Eitan Levin, Amit Singer

## Abstract

Single-particle cryo-electron microscopy (cryo-EM) has recently joined X-ray crystallography and NMR spectroscopy as a high-resolution structural method for biological macromolecules. In a cryo-EM experiment, the microscope produces images called micrographs. Projections of the molecule of interest are embedded in the micrographs at unknown locations, and under unknown viewing directions. Standard imaging techniques first locate these projections (detection) and then reconstruct the 3-D structure from them. Unfortunately, high noise levels hinder detection. When reliable detection is rendered impossible, the standard techniques fail. This is a problem especially for small molecules, which can be particularly hard to detect. In this paper, we propose a radically different approach: we contend that the structure could, in principle, be reconstructed directly from the micrographs, without intermediate detection. As a result, even small molecules should be within reach for cryo-EM. To support this claim, we setup a simplified mathematical model and demonstrate how our autocorrelation analysis technique allows to go directly from the micrographs to the sought signals. This involves only one pass over the micrographs, which is desirable for large experiments. We show numerical results and discuss challenges that lay ahead to turn this proof-of-concept into a competitive alternative to state-of-the-art algorithms.

## 1 Introduction

Cryo-electron microscopy (cryo-EM) is an imaging technique in structural biology used for single particle reconstruction (SPR) of macromolecules. In a cryo-EM experiment, biological samples are rapidly frozen in a thin layer of vitreous ice. The microscope produces a 2-D tomographic image of the samples embedded in the ice, called a *micrograph*. Each micrograph contains tomographic projections of the samples at unknown locations and under unknown viewing directions. The goal is to construct 3-D models of the molecular structure from the micrographs.

The signal to noise ratio (SNR) of the projections in the micrographs is a function of two dominating factors. On the one hand, the SNR is a function of the electron dose. To keep radiation damage within acceptable bounds, the dose must be kept low, which leads to high noise levels. On the other hand, the SNR is a function of the molecule size. The smaller the molecules, the fewer detected electrons carry information about them.

**Figure 1:**
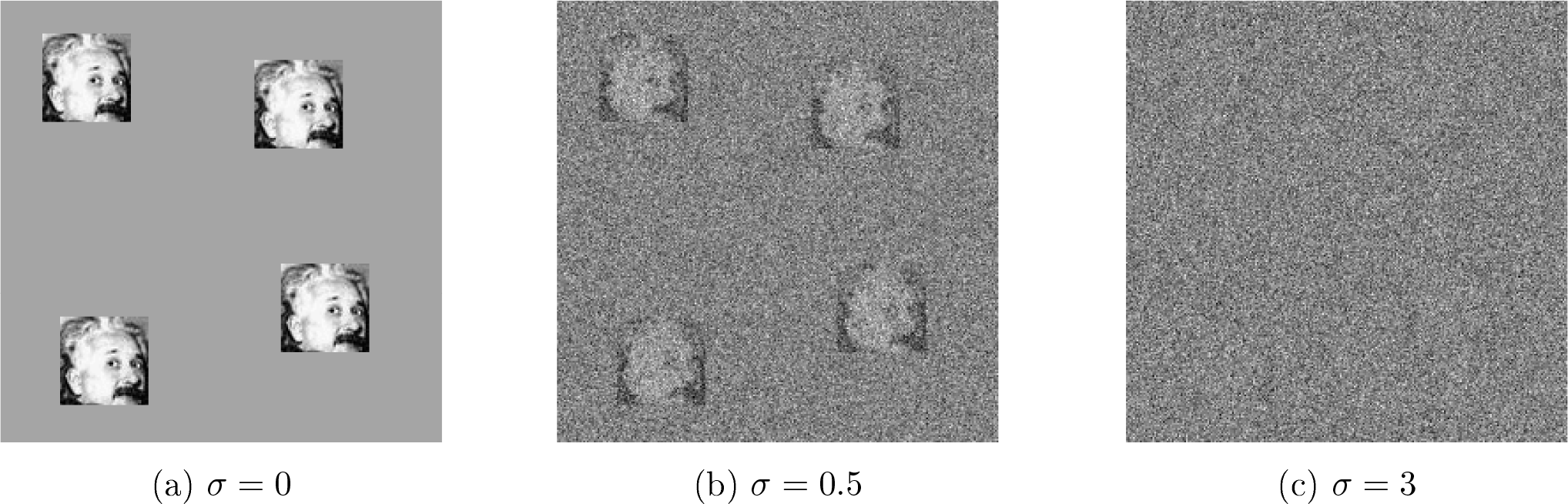
Example of micrographs of size 250 × 250 with additive white Gaussian noise of variance *σ*^2^ for increasing values of *σ*. Each micrograph contains the same four occurrences of a 50 × 50 image of Einstein. In panel (c), the noise level is such that it is very challenging to locate the occurrences of the planted image. In fact, it can be shown that at low SNR, reliable detection of individual image occurrences is impossible, even if the true image is known. By analogy to cryo-EM, this depicts a scenario where particle picking cannot be done.

**Figure 2:**
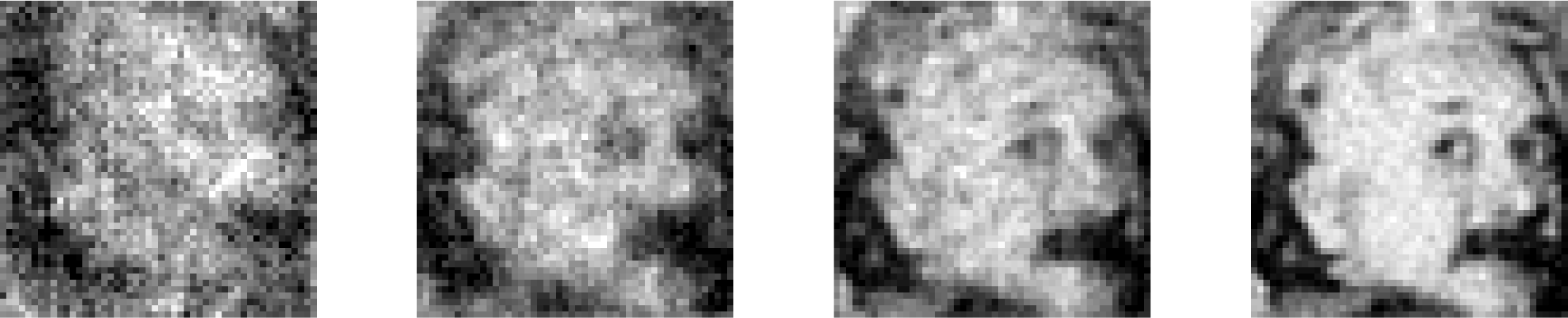
Recovery of Einstein from micrographs at noise level *σ* = 3 (see Figure 1(c)). Averaged autocorrelations of the micrographs allow to estimate the power spectrum of the target image. This does not require particle picking. A phase retrieval algorithm (RRR) produces the estimates here shown, initialized randomly. Estimates are obtained from 2 × 10^2^, 2 × 10^3^, 2 × 10^4^, 2 × 10^5^ micrographs (growing across panels from left to right) of size 4096 × 4096, each containing 700 image occurrences on average.

All contemporary methods in the field split the reconstruction procedure into two main stages. The first stage consists in extracting the various particle projections from the micrographs. This stage is called *particle picking*. The second stage aims to construct a 3-D model of the molecular structure from these projections. The quality of the reconstruction eventually hinges on the quality of the particle picking stage. Figure 1 illustrates how particle picking becomes increasingly challenging as the SNR degrades.

Crucially, it can be shown that reliable detection of individual particles is impossible below a certain critical SNR. This fact has been recognized early on by the cryo-EM community. In particular, in an influential paper from 1995, Richard Henderson [18] investigates the following questions:

> *For the purposes of this review, I would like to ask the question: what is the smallest size of free-standing molecule whose structure can in principle be determined by phase-contrast electron microscopy? Given what has already been demonstrated in published work, this reduces to the question: what is the smallest size of molecule for which it is possible to determine from images of unstained molecules the five parameters needed to define accurately its orientation (three parameters) and position (two parameters) so that averaging can be performed?*

In that paper and in others that followed (e.g., [15]), it was established that particle picking is impossible for molecules below a certain weight (below ~50 kDa). Joachim Frank voices a similar observation in his 2017 Nobel prize lecture: “*Using the ribosome as an example, it became clear from the formula we obtained that the single-particle approach to structure research was indeed feasible for molecules of sufficient size: Particle Size > 3/[Contrast^2^ × Resolution (as length) × Critical Electron Dose/*” [13]. As these two leaders of the cryo-EM community point out, it is impossible to reconstruct small molecules by any of the existing computational pipelines for single particle analysis in cryo-EM, because the particles themselves cannot be picked from the micrographs. Yet, imaging such molecules is important for structure-guided drug design and other applications [34].

The unique issues raised by small particles have been mitigated by recent technological advances in the field, including the use of Volta phase plates [22, 27] and scaffolding cages [29]. Despite this progress, detecting small molecules in the micrographs remains a challenge. We note that nuclear magnetic resonance (NMR) spectroscopy and X-ray crystallography are well suited to reconstruct small molecules. Yet, cryo-EM has a lot to offer even for molecules with already known structures obtained via NMR spectroscopy or X-ray crystallography, because these methods have limited ability to distinguish conformational variability.

In this paper, we argue that there is a gap between the two questions in Henderson’s quoted excerpt above, and that one may be able to exploit it to design better reconstruction algorithms. Specifically, the impossibility of particle picking does not necessarily imply impossibility of particle reconstruction. Indeed, the aim is only to reconstruct the molecule: estimating the locations of the particles in the micrograph is merely a helpful intermediate stage when it can be done. Our main message is that the limits particle picking imposes on molecule size do not necessarily translate into limits on particle reconstruction.

As a proof of concept, we first study a toy model for which it is easier to convey the mathematical principles. In this model, an unknown image appears multiple times at unknown locations in each of several micrographs, each affected by additive Gaussian noise-see Figure 1 for an illustration. The goal is to estimate the planted image. The task is challenging in particular when the SNR is low enough that particle picking (identifying the locations of each image in each micrograph) cannot be done reliably. This problem is interesting on its own as it appears in other scientific applications, including spike sorting [26], passive radar [16] and system identification [30].

In order to recover the image, we use autocorrelation analysis. Specifically, we relate the autocorrelations of the micrographs to the autocorrelations of the image. For any noise level, these autocorrelations can be estimated to any desired accuracy, provided that we observe sufficiently many image occurrences and the latter are separated in the micrograph. Importantly, there is no need to detect individual image occurrences. The autocorrelations of the micrographs are straightforward to compute and require only one pass over the data. After estimation of the density of particles in the micrographs, these directly yield estimates for the autocorrelations of the target image itself. To estimate the image itself from its estimated autocorrelations, we solve a nonlinear inverse problem; see for instance Figure 2.

Beyond this 2-D proof of concept, we look toward 3-D reconstruction as well. Zvi Kam [21] first proposed autocorrelation analysis for 3-D reconstruction, under the assumption of perfect particle picking: his method used autocorrelations of the picked, perfectly centered, particles. In contrast, we derive the mathematical relation between the autocorrelations of the micrographs as a whole and the 3-D volume, under some simplifying conditions. We show a few numerical examples and outline the future developments required to make this method applicable to experimental data.

Another interesting feature of the described approach pertains to model bias, whose importance in cryo-EM was stressed by a number of authors [37, 43, 19, 42]. In the classical “Einstein from noise’ experiment, multiple realizations of pure noise are aligned to a picture of Einstein using template matching and then averaged. In [37], it was shown that the averaged noise rapidly becomes remarkably similar to the Einstein template. In the context of cryo-EM, this experiment exemplifies how prior assumptions about the particles may influence the reconstructed structure. This model bias is common to all particle picking methods based on template matching. In our approach, no templates or human intervention are required, thus significantly reducing concerns about model bias.

## 2 Proof of concept: A mathematical toy model

In this section, we present a toy model in order to introduce the mathematical principles enabling estimation of a signal in a low SNR regime, even when detection is impossible. Later, we discuss how these principles carry through for the SPR problem. We first formulate the model for 1-D signals to ease exposition.

Let *x* ∈ ℝ^*L*^ be the target signal and let *y* ∈ ℝ^*N*^ be the observed data, where we assume *N* is far larger than *L*. Let *s* ∈ {0, 1}^*N* − *L*+1^ be a binary signal indicating (with 1’s) the starting positions of all occurrences of *x* in *y*, so that, with additive white Gaussian noise:

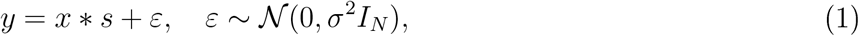

where * denotes linear convolution. While both *x* and *s* are unknown, the goal is only to estimate *x* from *y*. This is a key difference with other works on *blind deconvolution*, a longstanding problem arising in a variety of engineering and scientific applications such as astronomy, communication, image deblurring, system identification and optics; see [20, 36, 3, 2], just to name a few. The parameters of the signal *s* (the locations of its nonzero values) are the *nuisance variables* of the problem. As will be shown next, in a low SNR environment, estimating *s* is impossible, whereas estimating *x* is tractable under some conditions on *s*.

We assume the binary signal *s* obeys a *separation condition*:

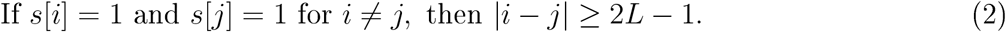

In words: the starting positions of any two occurrences must be separated by at least 2*L* − 1 positions, so that their end points are necessarily separated by at least *L* − 1 signal-free entries in the micrograph.

Our problem can be interpreted as a special case of the *system identification* problem. Similarly to (1), the forward model takes the form *y* = *x* * *w* + *ε* where *x* is the unknown signal (the system’s impulse response), *w* is an unknown (often random) input sequence, and *ε* is an additive noise. The goal of this problem is to estimate *x*, usually referred to as “identifying the system.’ The question of identifiability of *x* under this observation model is addressed for certain Gaussian and non-Gaussian *w* in [6, 23]. In the special case where *w* is binary and satisfies our separation condition, we recover our model.

Likelihood-based methods estimate *x* as the maximizer of some function *f*(*x* | *y*), where *f* is derived from the likelihood function of *x* given the observed signal *y*. If some prior is assumed on *x*, then *f*(*x* | *y*) can be taken to be the posterior distribution of *x* given the data; this is the simplest form of Bayesian inference. Methods based on such formulations are popular nowadays in cryo-EM; see for instance [38, 35]. Optimizing the function *f*(*x* | *y*) exactly is often intractable, and thus heuristic methods are used instead. One proposed technique is to use Markov Chain Monte Carlo (MCMC) [10]. In special cases, including the case where *w* is binary, expectation maximization (EM) has been used [10]. The EM method for discrete *w* is based on a certain “forward-backward’ procedure used in hidden Markov models [33]. However, the complexity of this procedure is superlinear in *N*, and therefore its usage is limited for big data sets.

Because likelihood methods are computationally expensive, methods based on recovery from moments have also been previously used for system identification. Methods based on the third-and fourth-order moments are described and analyzed in [28, 14, 41]. Building on such ideas, we focus on an autocorrelation analysis for (1).

## 3 The detection limit

In the low SNR regime—even if *x* is known-estimating the binary sparse signal *s* is impossible, that is, one cannot reliably detect occurrences of *x* in the micrograph *y*. To support this claim, we consider a strictly simpler problem: suppose an oracle identifies for us one interval of length *L* in the micrograph that either contains a full signal occurrence (plus noise), or contains just noise. Our task is to determine which one it is, that is, to determine whether the corresponding entry of *s* is 0 or 1. The oracle further provides the signal *x*, the probability *q* that the interval contains signal, and the noise variance *σ*^2^.

This decision problem can be abstracted as follows: We have two known vectors *θ*_0_ = *x* and *θ*_1_ = 0 in ℝ^*L*^. There is a random variable *η* taking values 0 or 1 with probabilities *q* and 1 − *q*, respectively. We observe a random vector *X* ∈ ℝ^*L*^ (an extract of the micrograph) with the following distribution: if *η* = 0, then 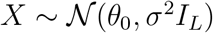 and if *η* = 1, then 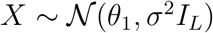.

We observe *X*, and our task is to estimate *η*. How reliably can this be done? If *q* ≥ 1/2, the constant estimator 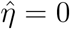 is correct with probability *q*; likewise, if *q* ≤ 1/2, the constant estimator 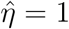 is correct with probability 1 − *q*. The question is, can we do better than this? We prove that, as *σ* → ∞, the answer is no. The result is proved in Appendix B.

### Proposition 3.1.

*For any deterministic estimator* 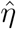 of *η*,

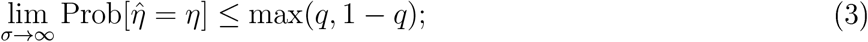

*that is: as the SNR deteriorates, the probability of success is no better than random chance*.

Proposition 3.1 implies that in order to estimate the signal at low SNR we must consider methods that aim to estimate the signal *x* directly, without estimating the nuisance variable *s* as an intermediate step. In the next sections, we consider autocorrelation analysis for that purpose.

## 4 Autocorrelation analysis

In general, for a signal *z* of length *m*, the autocorrelation of order *q* = 1,2, … is given for any integer shifts *l*_1_, …, *l*_*q*−1_ by

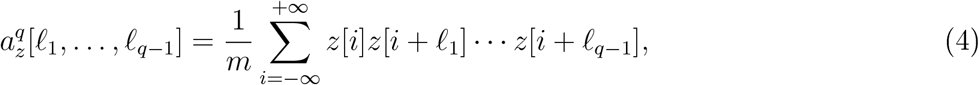

where indexing of *z* out of the range 0, …, *m* − 1 is zero-padded. Explicitly, the first-, second-and third-order autocorrelations are given by:

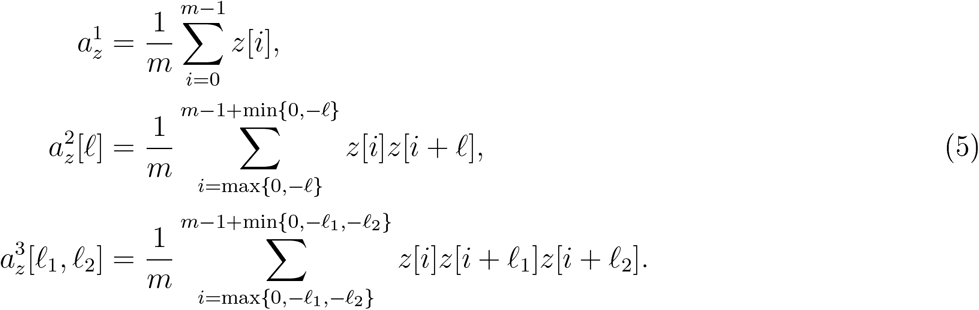

The autocorrelation functions have symmetries. Specifically, 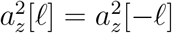, and 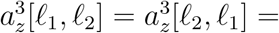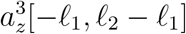 For our purposes, this will be applied both to *x* (of length *L*) and to *y* (of length *N*).

Under the separation condition, the relation between autocorrelations of the micrograph and those of *x* is particularly simple, as we now show. It is useful to introduce some notation: let *M* denote the number of occurrences of *x* in *y* (that is, the number of 1’s in *s*), and let

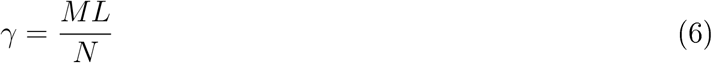

denote the density of *x* in *y* (that is, the fraction of entries of *y* occupied by occurrences of *x*.) The separation condition imposes 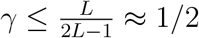.

For shifts in 0, …, *L* − 1, the autocorrelation functions of *y* depend on the corresponding autocorrelations of *x*, the noise level *σ* and the support signal *s*. Importantly, under the separation condition (2), the dependency on *s* is only through the density *γ*. We consider the asymptotic regime where *γ* remains constant; that is, as *N* goes to infinity, *M* also goes to infinity at the same rate (in other words, as we see an increasingly large micrograph, it contains increasingly many signal occurrences, with constant signal density). In that regime, the law of large numbers can be used to show the following statement:

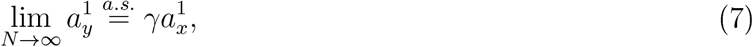

where equality holds almost surely (a.s.), meaning it holds with probability one. The randomness is over the Gaussian noise *ε*; *s* may be deterministic. Thus, given enough data, if *γ* is known, we can estimate 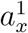 from *y*. (Ve show later how to estimate *γ* as well.)

We have a similar observation for the second-order autocorrelation: 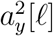 computes the correlation between *y* and a copy of *y* shifted by *l* entries. Considering *l* only in the range 0, …, *L* − 1, one can see that any given occurrence of *x* in *y* is only ever correlated with itself, and never with another occurrence. As a result,

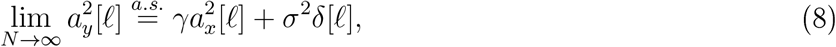

for *l* = 0, …, *L* − 1, where *δ*[*l*] equals one for *l* = 0 and zero otherwise. The last part captures the autocorrelation of the noise. Notice that, even if *σ* is unknown, entries *l* = 1, …, *L* − 1 still provide useful information about 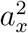.

Among the same lines, one can establish a relation for third-order autocorrelations:

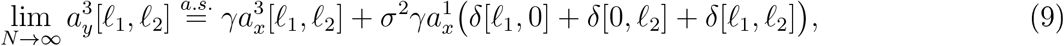

for *l*_1_, *l*_2_ = 0, …, *L* − 1, where *δ*[*l*_1_, *l*_2_] := *δ*[*l*_1_ − *l*_2_]. Here too, few entries are affected by *σ* in the limit. See Appendix C.

Computing the autocorrelations of the micrograph is straightforward. The natural question, treated next, is whether one can recover *x* from them.

## 5 Recovering a signal fron autocorrelations

A one-dimensional signal is determined uniquely by its second-and third-order autocorrelations. Indeed, assuming *z*[0] and *z*[*L* − 1] are nonzero (otherwise, redefine the length of the signal), we can recover *z* explicitly using this identity for *k* = 0, …,*L* − 1:

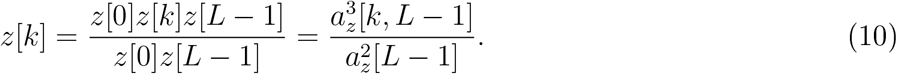

This proves the following useful fact:

#### Proposition 5.1

*A signal *z* ∈ ℝ^*L*^ is determined uniquely from* 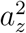 *and* 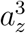.

A couple of remarks are in order. First, (10) is not numerically stable: if *z*[0] or *z*[*L* − 1] are close to 0, recovery of *z* is sensitive to errors in the autocorrelations. In practice, we recover *z* by fitting it to its autocorrelations using a nonconvex least-squares (LS) procedure, which is empirically more robust to additive noise; we have observed similar phenomena for related problems [5, 9, 1]. Second, note that the second-order autocorrelation is not by itself sufficient to determine the signal uniquely. However, for dimensions greater than one ^1^, almost all signals are determined uniquely up to sign (or phase for complex signals) and reflection through the origin (with conjugation in the complex case) [17]. The sign ambiguity can be resolved by the mean of the signal if it is not zero. However, determining the reflection symmetry still requires additional information, beyond the second-order autocorrelation.

The observed moments 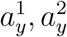 and 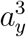 of *y* do not immediately yield the moments of the signal *x*, as seen by (7), (8) and (9); rather, the two are related by the noise level *σ* and the ratio *γ*. We will show, however, that *x* is still determined by the observed moments of *y*.

First, we observe that if the noise level *σ* is known, generally, one can estimate *γ* from the first two moments of the micrograph. The proof is provided in Appendix D.

### Proposition 5.2

*Let *σ* > 0 be fixed and assume that the separation condition (2) holds. If the mean of *x* is nonzero, then*

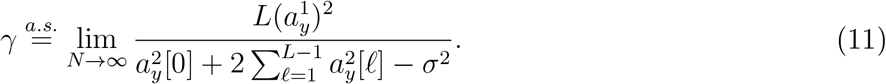

Using third-order autocorrelation information of *y*, both the ratio *γ* and the noise *σ* are determined. For the following results, when we say that a result holds for a “generic’ signal *x*, we mean that the set of signals which cannot be determined by these measurements has Lebesgue measure zero. In particular, this means that we can recover almost all signals with the given measurements. The proof is provided in Appendix E.

### Proposition 5.3.

*Assume *L* ≥ 3 and assume that the separation condition (2) holds. In the limit of *N* → ∞, the observed autocorrelations 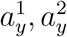 and 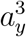 determine the ratio *γ* and noise level *σ* uniquely for a generic signal *x*. If 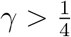, then this holds for any signal *x* with nonzero mean*.

From Propositions 5.1 and 5.3 we deduce the following:

#### Corollary 5.4.

*In the limit of *N* → ∞ and under the separation condition (2), the signal *x*, the ratio *γ*, and the noise level *σ* are determined from the first three autocorrelation functions of *y* if either the signal *x* is generic or *x* has nonzero mean and 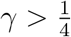*.

As a side note, under the separation condition, the length *L* of the signal can also be determined from the autocorrelations in the asymptotic regime, by inspection of the support of 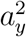.

## 6 Numerical experinents

The technique we advocate allows recovery of a signal hidden in noisy micrographs without detecting the location of the signals embedded in these micrographs. To illustrate the underlying principles of the method, we present several numerical examples for the toy model (1) and a simple proof of concept for simulated cryo-EM data. Appendix A provides additional details on the experiments. The code to generate all figures is publicly available in https://github.com/PrincetonUniversity/BreakingDetectionLimit.

### 6.1 Toy model

In the first experiment, we estimated an 50-by-50 pixel image of Einstein with mean zero from a growing number of micrographs, each of size 4096 × 4096 pixels. Each micrograph contains, on average, 700 occurrences of the target image at random locations. Thus, about 10% of each micrograph contains signal. The micrographs are contaminated with additive white Gaussian noise with standard deviation *σ* = 3, corresponding to 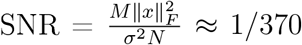. This high noise level is illustrated in Figure 1. To simplify the experiment, we assume the number of signal occurrences and the noise standard deviation are known. Micrographs are generated such that any two occurrences are always separated by at least 49 pixels in each direction in accordance with the separation condition (2).

We compute the average second-order autocorrelation of the micrographs. This is a particularly simple computation which can be efficiently executed with a fast Fourier transform (FFT) in parallel. Given the noise level and number of image repetitions, the second-order autocorrelation of the image can be easily deduced from (8). Then, to estimate the target image, we resort to a standard phase retrieval algorithm called relaxed-reflect-reflect (RRR) [12], initialized randomly. Relative error is measured as the ratio of the root mean square error to the norm of the ground truth (square root of the sum of squared pixel intensities).

Figure 2 shows several estimated images for a growing number of micrographs. Figure 3 presents the normalized recovery error as a function of the amount of data available. This is computed after fixing the reflection symmetries (see Section 4). As evidenced by these figures, the ground truth image can be estimated increasingly well from increasingly many micrographs, without particle picking.

**Figure 3:**
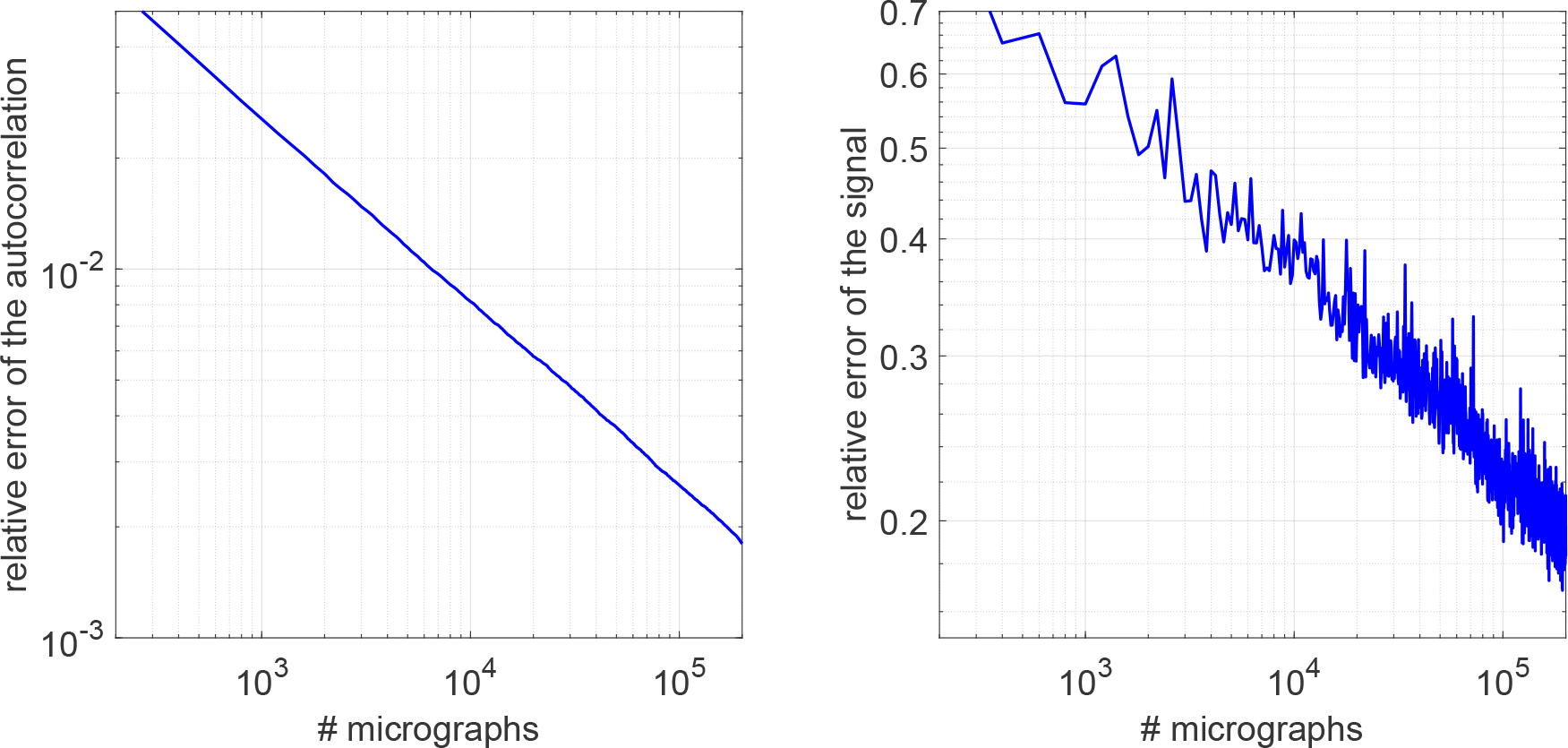
Relative error curves for the experiment of Figure 2.

In practice, we do not expect to know *γ* and maybe not even *σ*. Figure 4 shows recovery of a 1-D signal from the first three autocorrelations of the data. The autocorrelations are computed from noisy micrographs with *σ* = 3 that obey the separation condition of (2); see Figure 5. Both the signal and *γ* are estimated simultaneously from the observed autocorrelations by LS fitting; more details are given in Appendix A. Crucially, the LS does not require knowledge of *σ*.

### 6.2 Cryo-EM

The application of autocorrelation analysis to cryo-EM follows the same mathematical principles. The derivation of the first three autocorrelations of the micrographs and their relations to the volume itself are provided in Appendix F. In particular, numerical evidence suggests that the third-order autocorre-lation uniquely determines the 3-D volume. Figure 6 shows recovery of the 3-D volume from the clean autocorrelations and from noisy micrographs. Excerpts from the noisy micrographs are shown in Figure 7. Unfortunately, the mapping between the autocorrelations and volume seems to be ill-conditioned, preventing high-resolution recovery from noisy data. The details of the reconstruction algorithm are given in Appendix A. In the next section, we outline how we suggest to overcome the ill-conditioning in future work.

**Figure 4:**
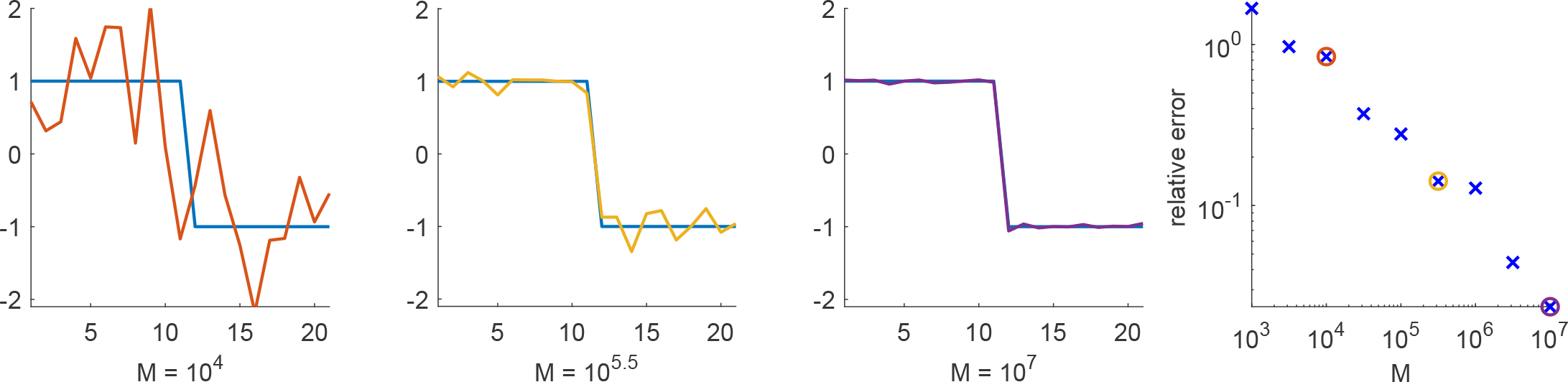
Recovery of a 1-D signal of length *L* = 21 at noise level *σ* = 3 from *M* repetitions. The length of the micrograph was set to be *N* = 410 * *M*. The first three panels (left to right) show reconstruction with different *M* values compared to the ground truth signal (in blue). The true value of *γ* is 0.0512. The relative errors of *γ* for the three panels are: 4.8%, 4%, 1.2%. The right panel shows the relative error as a function of *M*.

**Figure 5:**
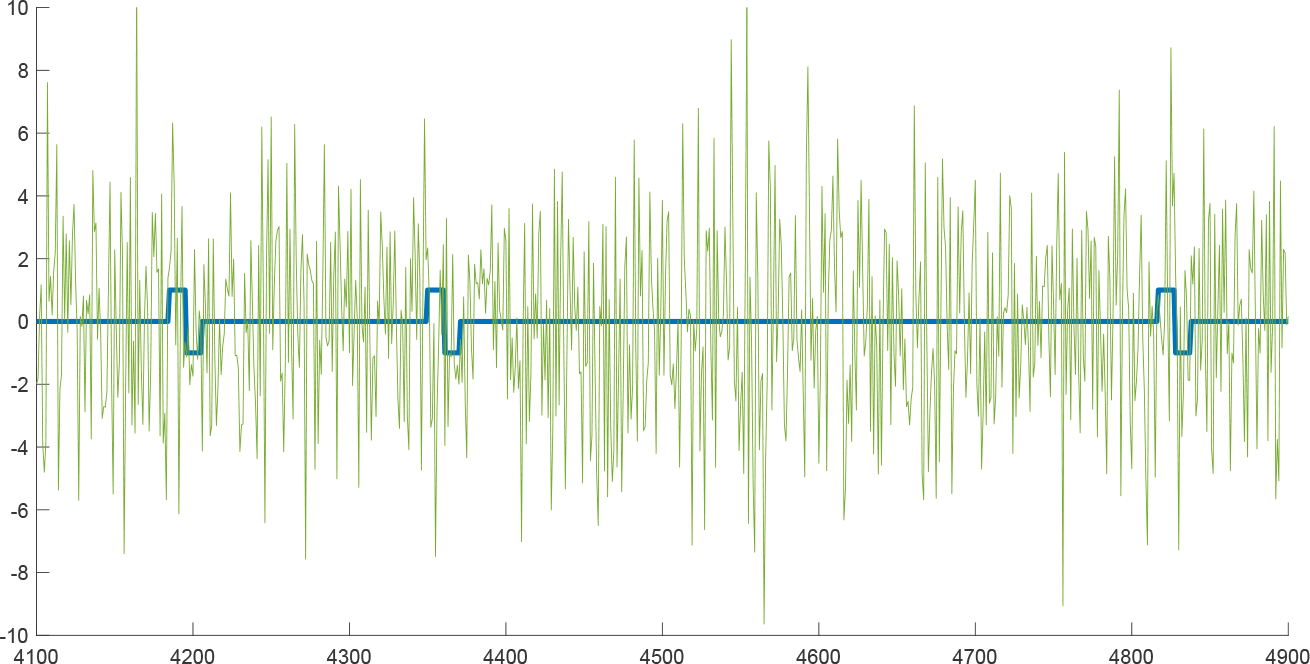
An example for a 1-D noisy measurement for the experiment in Figure 4. The signal occurrences appear in blue, and the noisy data in green.

While we cannot provide a high-resolution 3-D reconstruction from noisy data with the current algorithm, our method can be easily applied to the problem of deciding whether a micrograph contains projections or merely pure noise—a problem considered in classical works in statistics [11] and cryo-EM [19]. This task can be performed by considering solely the recovered *γ* (the fraction of pixels occupied by projections in the micrograph), estimated as part of the recovery algorithm. Figure 9 presents excerpts of two noisy micrographs, only one of which contains projections. In the presence of projections, the estimated *γ* was 0.12, corresponding to approximately 6784 projections. On the other hand, the estimated *γ* drops to 10^−5^ for the pure noise micrograph, corresponding to less than one projection.

**Figure 6:**
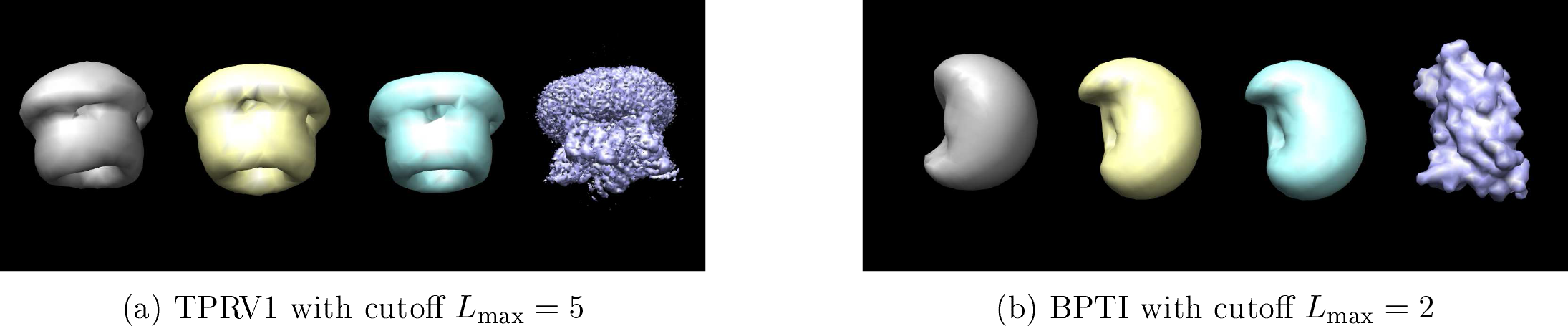
Reconstructions from the first, second and third order autocorrelations. The ground truth volumes were expanded according to (25) with cutoff *L*_max_. The original molecules are shown in purple and the (smoothed) ground truths in blue to illustrate the smoothing effect of our downsampling and truncation of the spherical harmonics expansion. The reconstructions from clean autocorrelations are shown in yellow and recoveries from autocorrelations estimated from noisy data in gray. For the noisy experiments, we used 300 micrographs with SNRs of 1/16 for TRPV1 and 1/64 for BPTI. We present excerpts from noisy micrographs side-by-side with the corresponding clean ones in Figure 7 (see appendix).

**Figure 7:**
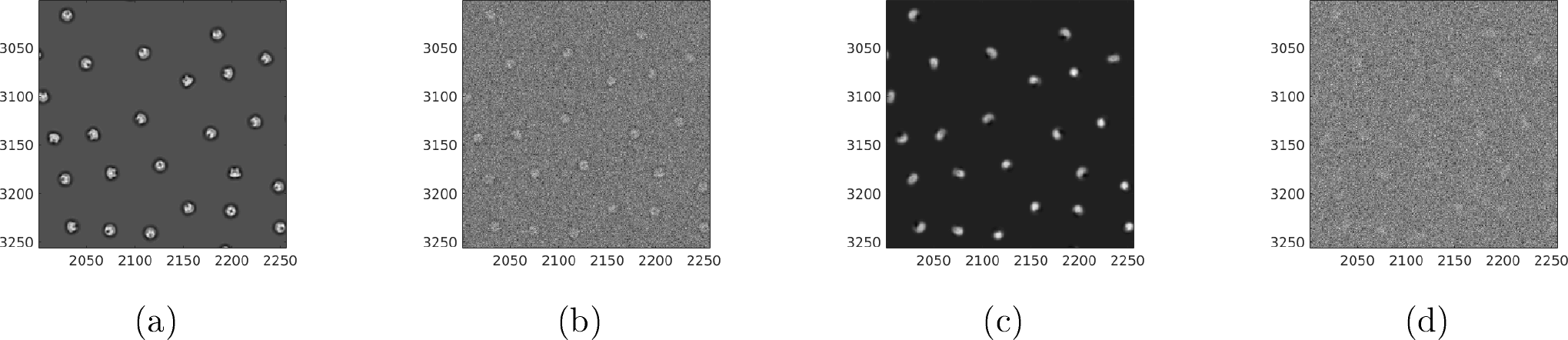
Excerpts from the noisy micrographs used for the reconstructions in Figure 6 and the corresponding clean excerpts. (a) Excerpt of the clean TRPV1 micrograph; (b) Excerpt of the noisy TRPV1 micrograph with SNR = 1/16; (c) Excerpt from the clean BPTI micrograph; (d) Excerpt from the noisy BPTI micrograph with SNR = 1/64.

**Figure 8:**
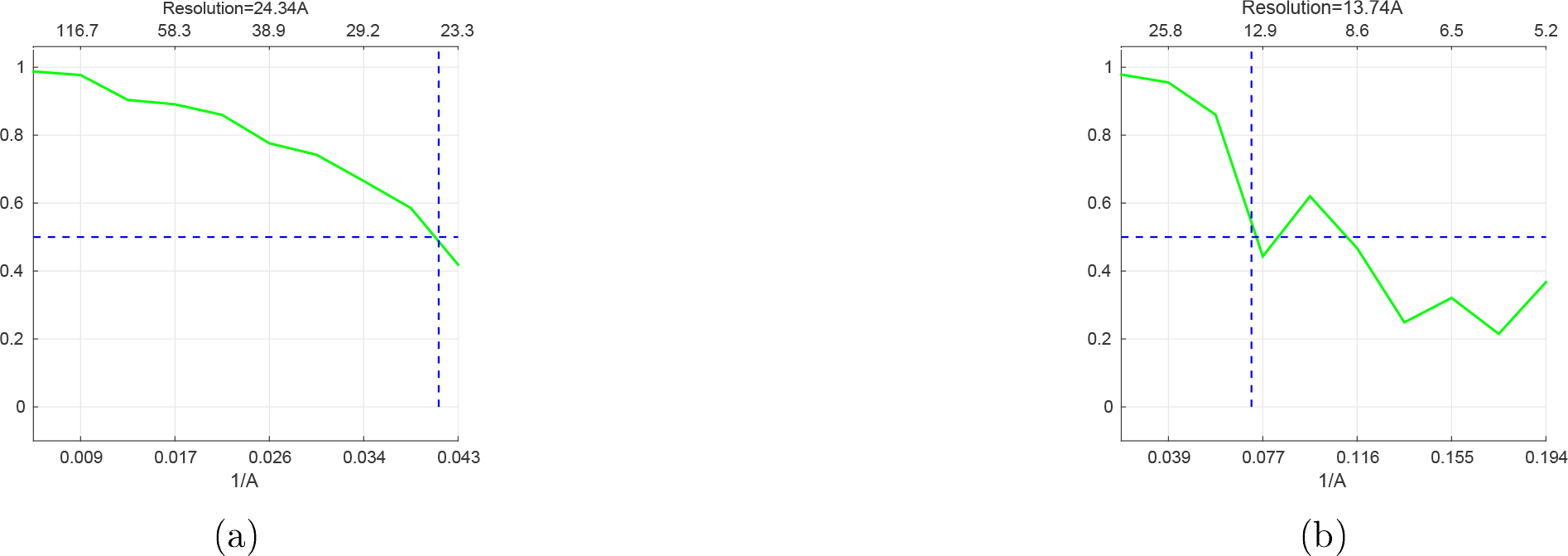
Fourier Shell Correlations (FSCs) for the noisy reconstructions presented in Figure 6 using the 0.5 criterion, compared to the (smoothed) ground truth. (a) FSC for the TRPV1 reconstruction, giving resolution of 24Å; (b) FSC for the BPTI reconstruction, giving resolution of 13Å.

## 7 Discussion

In the simplified mathematical model above, we showed it is possible to estimate a signal without detecting its appearances. Our strategy is to compute autocorrelations of the micrographs and to relate these statistics to the unknown signal’s parameters. Recovering the parameters from the statistics reduces to solving a set of polynomial equations.

**Figure 9:**
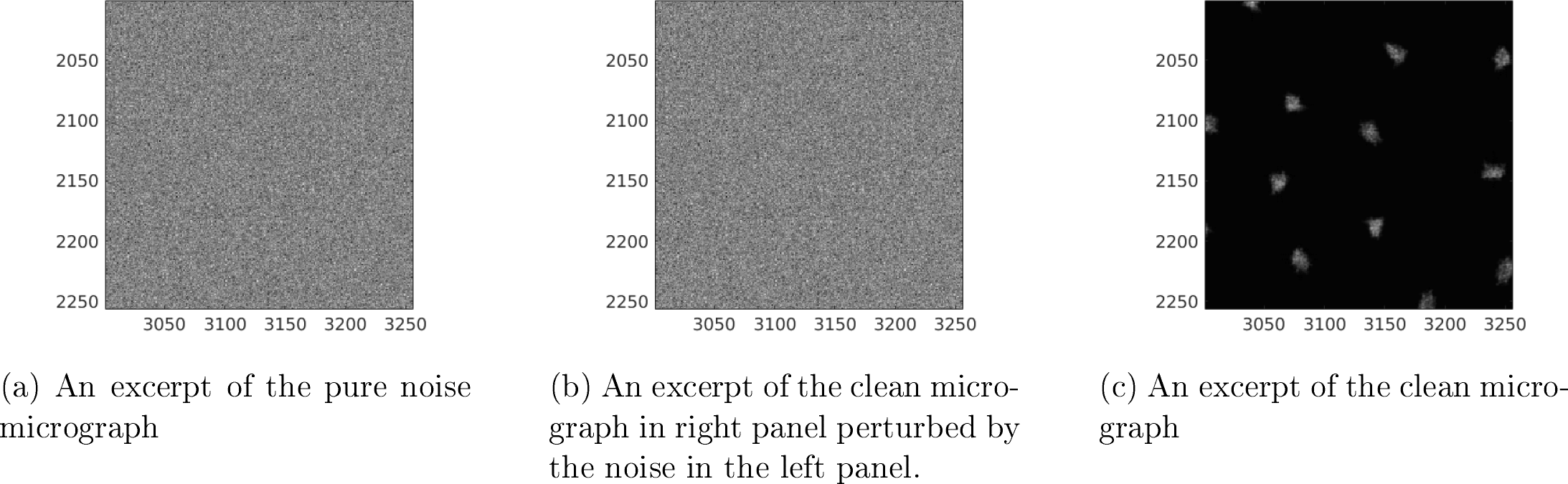
All micrographs are of size 7420^2^ pixels and the projections are taken from the BPTI molecule of size 31^3^. The added noise was drawn from i.i.d. Gaussian distribution with zero mean and standard deviation 25, corresponding to an SNR below 1/1024. The noise realization is identical in both micrographs.

We also showed how this technique can, in principle, be applied to cryo-EM. Crucially, the outlined approach involves no particle picking, hence a fortiori no viewing direction estimation. Concerns for model bias are also greatly reduced since no template matching is involved. Additionally, our technique also allows the use of much lower defocus values. Lower defocus means lower contrast, but will maintain higher frequency information. Consequently, we may be able to get high resolution reconstructions from fewer micrographs.

Looking toward applying the framework to encompass all important features of real cryo-EM experiments, our work implies that it might be possible to reconstruct small molecules, particularly, molecules that are too small to be detected in micrographs. In pursuing this research direction, our goal is to significantly increase the range of molecules to which cryo-EM can be successfully applied. We recognize that significant challenges lay ahead for the implementation of the proposed approach to high-resolution 3-D reconstruction directly from the micrographs. We discuss a few now.

The numerical experiments we have performed reveal that the third-order autocorrelation may not be enough for 3-D reconstruction in practice, due to high sensitivity. This suggests that fourth-order autocorrelation may be necessary. This in turn would imply that the procedure might require a large amount of data. Recent trends in high-throughput cryo-EM technology give hope that this may be a lesser concern in the long term. Still, large amounts of data also require large amounts of computation. On this front, we note that computing autocorrelations can be executed efficiently on CPUs and GPUs, and in parallel across micrographs. It can even be done in a streaming mode, as only one pass through each micrograph is necessary. The output of this data processing stage is a summary in the form of autocorrelation estimates: its size is a function of the resolution, not a function of the number of observed micrographs. Subsequent steps, which involve solving the system of polynomial equations, scale only in the size of that summary. Of course, an important question then is whether the equations can be solved accurately and efficiently in practice.

To reach high-resolution reconstruction, beyond data acquisition and computational challenges, there are modeling issues to consider. In contrast to the simplifying assumptions we have made above, the noise might be colored; the viewing directions of the particles may be distributed non-uniformly; there may be conformational heterogeneity; particles generally do not satisfy our separation condition; and micrographs undergo a contrast transfer function which we have omitted. All of these aspects can be handled with the same general strategy: establish a forward model relating the expected autocorrelations of the micrographs to the target volume(s) and all parameters necessary to model the above effects. For instance, for colored noise, the forward model may involve multiple parameters to capture the power spectrum of the noise instead of the single parameter *σ*^2^. Similarly, instead of the separation condition, we can model the spacing between the projections using a parameterized pair-correlation function. Such a function models the distribution of distances between neighboring projections. The observed autocorrelations depend linearly on these parameters, which would be estimated as part of the inverse problem. All these aspects must be taken into account so the method can be applied on experimental data. We hope to take care of these issues in future research.

## Acknowledgnent

The authors thank Ayelet Heimowitz, Joe Kileel, Roy Lederman, Amit Moscovich, Nir Sharon and Fred Sigworth for helpful discussions, and Boris Landa and Yoel Shkolnisky for providing the code for the 2-D PSVFs expansion. The research was partially supported by Award Number R01GM090200 from the NIGMS, FA9550-17-1-0291 from AFOSR, Simons Foundation Math+X Investigator Award, and the Moore Foundation Data-Driven Discovery Investigator Award. NB is partially supported by NSF award DMS-1719558.

## A Numerical experiments details

We run the toy model experiments on a shared computer with 144 logical CPUs of type Intel(R) Xeon(R) CPU E7-8880 v3 @ 2.30GHz and 792 Gb of RAM; we use at most 72 of these CPUs.

### 2-D experiment

For the 2-D experiment shown in Figures 2 and 3, we generated micrographs of size 4096 × 4096 pixels. In each micrograph, we placed Einstein’s image (of zero mean) of size 50 × 50 in random locations, while preserving the separation condition (2). This is done by randomly selecting 4000 placements in the micrograph, one at a time with an accept/reject rule based on the separation condition and locations picked so far. On average, 700 images are placed in each micrograph. Then, i.i.d. Gaussian noise with standard deviation *σ* = 3 is added, inducing an SNR of approximately 1/370. An example of a micrograph’s excerpt is presented in the right panel of Figure 1.

In this experiment, we assume to know the noise level *σ* and the total number of occurrences of the target image across all micrographs. In stark contrast with the 1-D setup, the second-order autocorrelation determines almost any target image uniquely, up to reflection through the origin. This is because the second-order autocorrelations correspond to the Fourier magnitudes of the signal through the 2-D Fourier transform. Therefore, we estimate the signal’s Fourier magnitudes (or power spectrum) from the Fourier magnitudes of the micrographs, at the cost of one 2-D fast Fourier transform (FFT) per micrograph. These can be computed highly efficiently and in parallel. Computing the power spectrum of 200 micrographs took less than 23 seconds.

To recover the target image from the estimated power spectrum, we use a standard phase retrieval algorithm called relaxed-reflect-reflect (RRR). This algorithm iterates the map

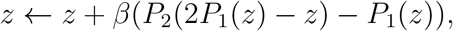

on an image *z* of size 2*L* × 2*L*. We set the parameter *β* to 1. The map is designed so that, if the estimated power spectrum is exact, then fixed points contain Einstein’s image in the upper-left corner of size *L* × *L*, possibly reflected through its origin, and zeros elsewhere. The operator *P*_2_(*z*) combines the Fourier phases of the current estimation *z* with the estimated Fourier magnitudes. The operator *P*_1_(*z*) zeros out all entries of *z* outside the *L* × *L* upper-left corner. In all experiments, the algorithm halted after a fixed number of 2000 iterations. Running the RRR took about 3 seconds.

The computational complexity scales as *O*(*N* ^2^*L*^2^ + *F* (*L*)), where *F* is the complexity of the RRR algorithm and *N*^2^ stands for the total number of pixels across micrographs.

### 1-D experiment

For the 1-D experiment depicted in Figure 4, we used a signal of length *L* = 21 and generated an observation *y* of length *N* = 10(2*L* − 1)*M*, where *M* is the number of repetitions. The observation was generated by randomly selecting placements in *y*, one at a time with an accept/reject rule based on the separation condition and locations picked so far until reaching the desired number of signal occurrences. Then, i.i.d. Gaussian noise with mean zero and standard deviation *σ* = 3 is added, to form the observed *y*. The resulting SNR of *y* is about 1/175. Figure 5 shows an example of the observed data.

Given the observation *y*, we proceed to compute the autocorrelations. We excluded biased elements of the autocorrelations; that is 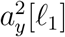 and 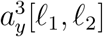 such that *l*_1_, *l*_2_ or *l*_1_ − *l*_2_ are zero. This has the practical effect that we need not know *σ*. By also taking symmetries into account, the third-order autocorrelation contains 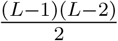 remaining entries. Thus, in total we have

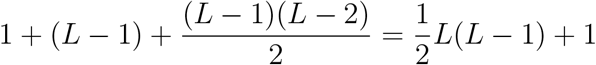

coefficients. This redundancy suggests that it might be possible to estimate several signals simultaneously (compare with [9, 4]). In practice, the autocorrelations are computed on disjoint segments of *y* of length 100·10^6^ (if the length of the measurement is larger than that) and added up, without correction for the junction points.

Having computed the moments of interest, we now estimate the signal *x* and coefficient *γ* which agree with the data. We choose to do so by running an optimization algorithm on the following nonlinear LS problem:

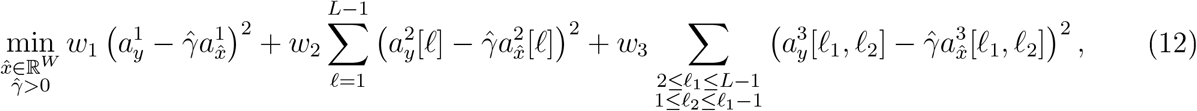

where *W* ≥ *L* is the length of the sought signal and the weights are set to *w*_1_ = 1/2, *w*_2_ = 1/2*n*_2_, *w*_3_ = 1/2*n*_3_, where *n*_2_, *n*_3_ are the number of moments used: *n*_2_ = *L* − 1, 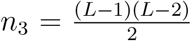 (weights could also be set in accordance with variance estimates as in [9]).

Setting *W* = *L* (as is a priori desired) is problematic because the above optimization problems appears to have numerous poor local optimizers. A similar phenomenon was recently observed in a related problem [44]. Thus, we first run the optimization with *W* = 2*L* − 1. This problem appears to have few poor local optima, perhaps because the additional degrees of freedom allow for more escape directions. Since we hope the signals estimated this way correspond to the true signals zero-padded to length *W*, we extract from each one a subsignal of length *L* that has largest *l*_2_-norm. This estimator is then used as initial iterate for (12), this time with *W = L*. We find that this procedure is reliable for a wide range of experimental parameters. To solve (12), we run the trust-region method implemented in Manopt [8], which allows to treat the positivity constraints on coefficients 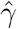. Notice that the cost function is a polynomial in the variables, so that it is straightforward to compute it and its derivatives.

The computational complexity scales as *O*(*NL*^2^ + *F* (*L*)), where *F* is the complexity of the solving the LS. For example, computing the moments for the experiment in the left panel of Figure 4 took 38 seconds, while the optimization took 5.5 seconds in total.

### 3-D experiments

The experiments in Figure 6, and the recoveries of *γ* for the experiment in Figure 9, were performed on a machine with 40 cores of Intel Xeon E5-2698 v4 @ 2.20GHz with 100 GB of RAM, and took 2 hours per reconstruction. The computation of the moments from each micrograph in Figure 9 was performed on a machine with 4 nVidia P100 GPUs with 16 GB of memory each and with 100 GB of RAM. It took 3 minutes per micrograph to compute the first three autocorrelations.

For the clean experiments in Figure 6, we computed the analytical first three autocorrelations of the volume as explained in Appendix F. To estimate the coefficients of the volume itself, we solve the LS problem

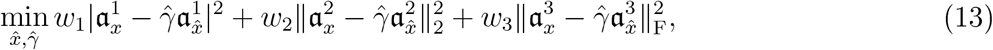

where the explicit expressions of 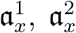 and 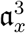 are given in (46), (45) and (44), respectively. In the experiments, we set *w*_1_ = *w*_2_ = *w*_3_ = 1. The LS problem was solved using Matlab’s `lsqnonlin` solver for nonlinear LS problems. The expected 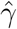 in this experiment is one.

The computational complexity for the computation of the moments from the micrograph is *O*(*N*^2^*P*^6^), since we have *N*^2^ patches that we extract from the micrograph. For each patch we perform an expansion in PSVFs with complexity *O*(*P*^3^) [24], and then compute the autocorrelations dominated by computation of the bispectrum as in (38) costing *O*(*P*^6^). The complexity of evaluating the bispecturm using (44) after the precomputation takes *O*(*BP*^6^), where *B* is the number of entries in the bispectrum. Since *B* = *O*(*P*^3^), the total complexity for evaluating the bispectrum is *O*(*P*^9^). That is because, if the number of volume expansion coefficients is *V* = *O*(*P*^3^), the dominant step can be written as a matrix-vector multiplication with a matrix of size *BV* × *V* and a vector *V* × 1, so the cost is *BV*^2^ = *O*(*P*^9^). Thus, the complexity of solving the LS problem (13) is *O*(*KP*^9^) where *K* is the number of iterations required for the optimizer to converge, since the bispectrum evaluation dominates the cost of each iteration.

The true volume used in the experiments in Figures 9 and 6b was the Bovine Pancreatic Trypsin Inhibitor (BPTI) mutant with altered binding loop sequence, whos atomic model is available in the Protein Data Bank (PDB) as 1QLQ^2^. We generated an EM map from this atomic model in UCSF Chimera [32] at a resolution of 5 Å, and cropped it to remove zeros at the boundary to obtain a volume of size 31^3^. For the experiment in Figure 6b, the volume was downsampled to size 20^3^. For the experiment in Figure 6a, we used the TRPV1 in complex with DkTx and RTX, whose EM map is available in the Electron Microscopy Data Bank (EMDB) as EMD-8117^3^. The original map has size 192^3^, and was downsampled to size 20^3^. To generate the ground truth for our reconstructions, we expanded both volumes as in (25) with cutoff *L*_max_ = 5 for TRPV1 and *L*_max_ = 2 for BPTI. For the TRPV1 reconstruction, the optimizer converged to a point giving relative *l*_2_ error of 10^−6^ in the first three autocorrelations and an error of 10^−1^ in the expansion coefficients of the volume. For the BPTI reconstruction, the errors in the autocorrelations were 10^−6^, 10^−7^ and 10^−8^ for the third, second and first autocorrelations, respectively, while the error in the expansion coefficients of the volume was 5 × 10^−2^. This illustrates the ill-conditioning of the map between the volume and its first three autocorrelations that prevents us from obtaining high-resolution results from noisy data.

The micrographs for the experiments presented in Figures 9 and 7 were generated as follows. We sample rotation matrices from SO(3) uniformly at random using the QR-based algorithm described in [40], and generate the projection of the volume corresponding to that rotation matrix using `cryo_project` in ASPIRE^4^. The projections for the experiments in Figure 7 were obtained from the smoothed volumes, not the original ones. We keep track of the indices at which the upper left corner of a projection can be placed without violating the separation condition, so all projections are separated by at least *L* − 1 pixels in each dimension, where the projections are contained in a box of size *L* × *L*. The location of the upper left corner of each new projection is picked uniformly at random from the set of available indices. We continue adding projections to the micrograph until no more projections can be added without violating the separation condition. In the experiments, we define SNR slightly differently than in the toy model, namely, as 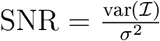 where 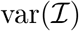 is the variance of our stack of micrographs and *σ*^2^ is the variance of noise. The noise level in the noisy micrograph in Figure 9 was SNR = 1/1024, the noise level in Figure 7 was SNR = 1/16 for the TRPV1 micrographs and SNR = 1/64 for the BPTI micrographs. We used 300 micrographs for each of the noisy reconstructions presented in Figure 6.

For the noisy reconstructions presented in Figure 6, the LS problem (13) was solved assuming spherical harmonic cutoffs of *L*_max_ = 5 for TRPV1 and *L*_max_ = 2 for BPTI, same as for the ground truth. For the experiment in Figure 9, the LS problem (13) was solved assuming the spherical harmonic cutoff for the volume is *L*_max_ = 0, which is sufficient to recover a significant 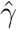 in the presence of projections in the micrograph.

### B Proof of Proposition 3.1

The proof is based on a variant of the Neyman-Pearson Lemma to derive the best (deterministic) estimator 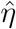. Take any estimator 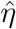; it is characterized by *S*: the set of *X*’s where 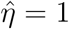, where *X* is a random vector as defined above Proposition 3.1. We write Prob_*i*_ to mean the probability conditional on the event *η* = *i*; that is, Prob_*i*_[*A*] = Prob[*A*|*η* = *i*]. Then, the probability that 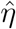 fails is:

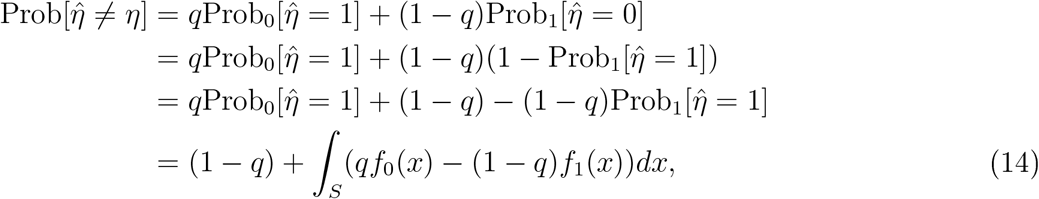

where *f*_*i*_(*x*) is the normal density with mean *θ*_*i*_ and variance *σ*^2^. The best estimator of *η* based on *X* minimizes the failure probability; hence, it minimizes the integral in (14) through an appropriate choice of the set *S*. This is achieved by picking all *x*’s such that the integrand is nonpositive:

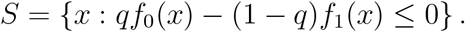

With Λ(*x*) = *f*_0_(*x*)/*f*_1_(*x*) and *b* = (1 − *q*)/*q*, the corresponding estimator is:

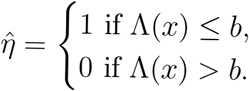

Taking logarithms, the set *S* can be rewritten as the set of *x*’s where:

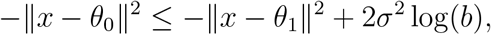

or equivalently

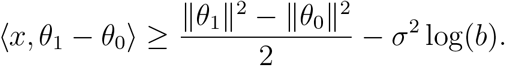

Now let us compute the probability of failure conditional on the event *η* = 0. In this case, failure occurs when *X* ∈ *S*. Since *X*|(*η* = 0) ∼ *N* (*θ*_0_, *σ*^2^), we can write *X*(*η* = 0) = *σZ* + *θ*_0_, where *Z* ~ *N*(0, *I*). On that condition,

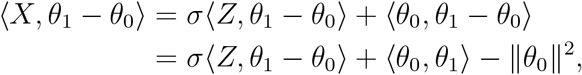

and failure occurs when

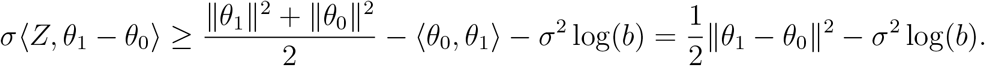

Define *Y* = 〈*Z*, *θ*_1_ − *θ*_0_〉 ~ *N*(0, ‖*θ*_1_ − *θ*_0_‖^2^) and divide through by *σ*. The above event is equivalent to:

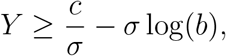

where *c* = ‖*θ*_1_ − *θ*_0_‖^2^/2. For simplicity, let us assume ‖*θ*_1_ − *θ*_0_‖ = 1, so that *Y* ~ *N*(0, 1). Then,

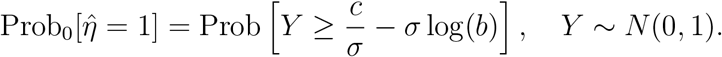

Similarly,

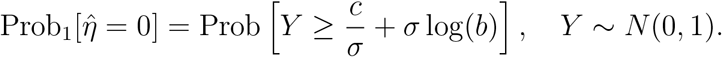

Thus, the overall probability of failure is:

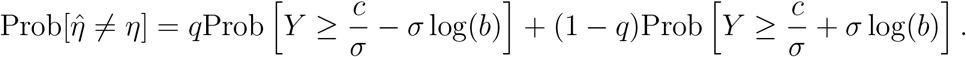

Now, if *q* = 1/2, then log(*b*) = 0. Hence the probability of failure is simply:

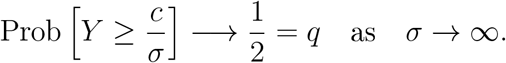

If *q* > 1/2, then *q* > 1- *q* and log(*b*) < 0. Consequently,

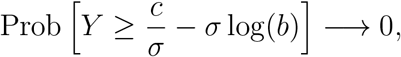

while

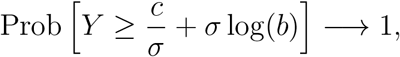

as *σ* → *∞*. Hence,

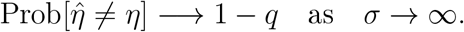

That is, the probability of success converges to *q*. Finally, if *q* < 1/2, then log(*b*) > 0 and a similar reasoning shows the probability of success converges to 1 − *q*. In all cases, the probability of success of the best possible deterministic estimator converges to max(*q*, 1- *q*).

### C Derivation of the identities in Section 4

We consider the asymptotic regime *N*, *M* → ∞ and assume that *M* = Ω(*N*), so that

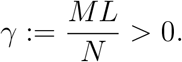

Any vectors with indices out of range are given value 0.

We start by considering the mean of the data:

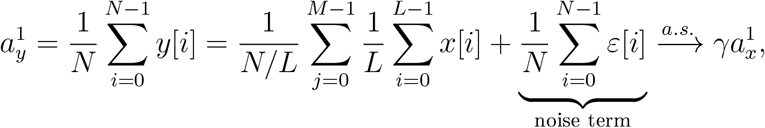

where the noise term converges to zero almost surely (a.s.) by the strong law of large numbers.

We proceed with the (second-order) autocorrelation for fixed *l* ∈ [0, …, *L* − 1]. We can compute:

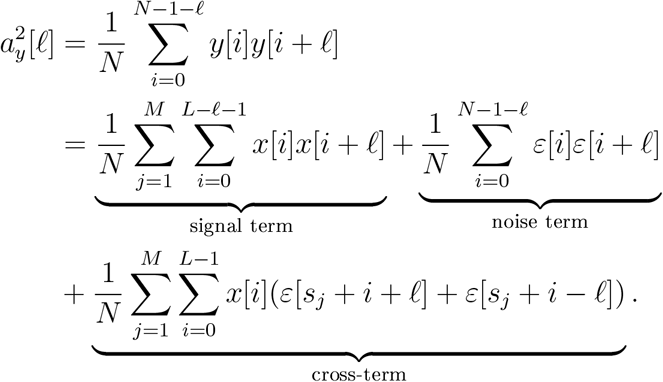

The cross-term is linear in the noise, and is easily shown to vanish almost surely in the limit *N* → ∞, by the strong law of large numbers. Ve break the signal term into *M* different sums, each containing one copy of the signal. This gives:

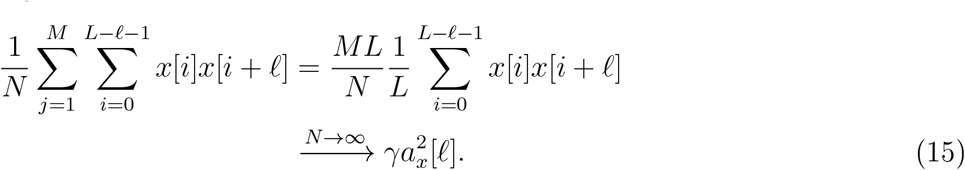

We next analyze the pure noise term. Vhen *l* = 0, we can break the noise term into a sum of independent terms:

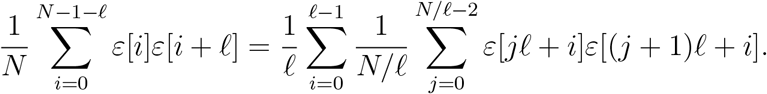

Each sum 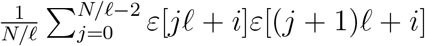 is an average of *N*/*l* independent terms with expectation zero, hence converges to zero almost surely as *N* → ∞. If *l* = 0, then we have:

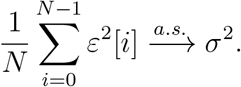

We now analyze the third-order autocorrelation. Let us *l*_1_ ≥ *l*_2_ ≥ 0. We have:

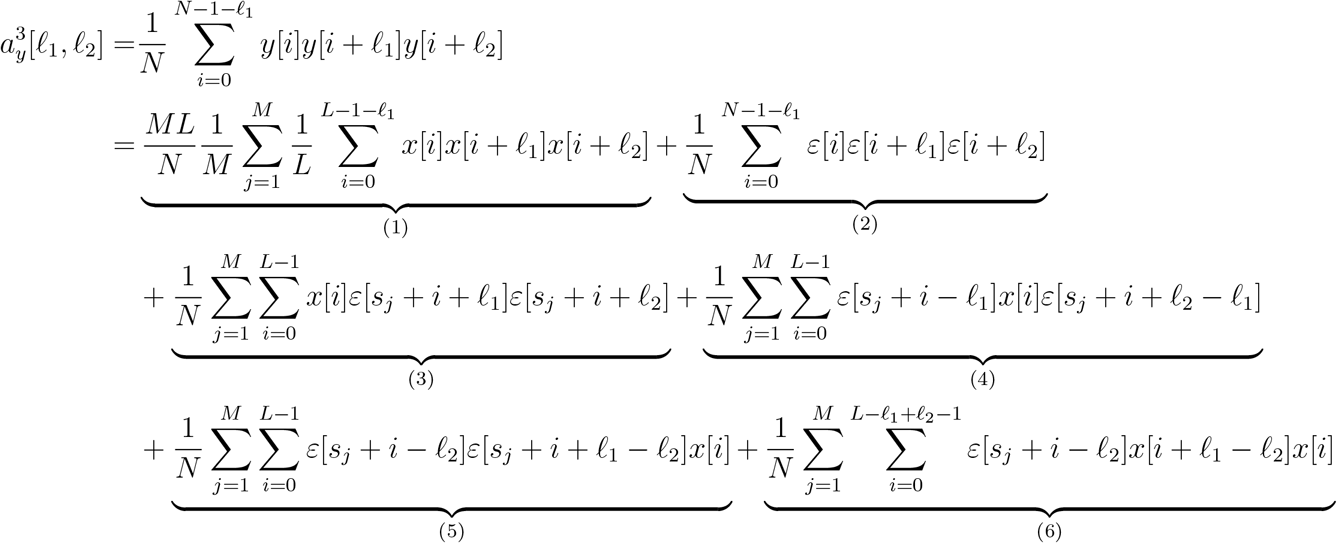

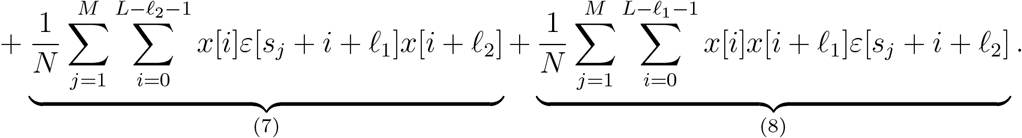

Terms (6), (7) and (8) are linear in *ε*, and can easily be shown to converge to 0 almost surely by the law of large numbers, by similar arguments as used previously. Term (1) converges to 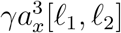 almost surely, for the same reasons as (15). To deal with terms (2)–(5), we must distinguish between different values of *l*_1_ and *l*_2_.

#### Case 1

0 < *l*_2_ < *l*_1_. Here, all summands with elements of *ε* involve products of distinct entries, which have expected value 0. Consequently, the usual argument shows that terms (2)–(5) all converge to 0 almost surely as *N* → ∞.

#### Case 2

0 = *l*_2_ < *l*_1_. Term (2) is an average of products of the form *ε*[*i*]^2^*ε*[*i* + *l*_1_], which have mean zero; consequently, term (2) converges to 0 almost surely. The same argument as for Case 1 shows that (3) and (5) also converge to 0. For term (4), we write:

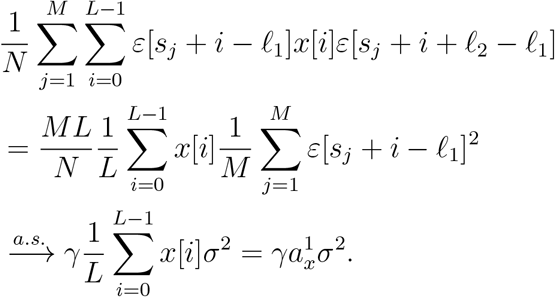

#### Case 3

0 < *l*_2_ = *l*_1_. An argument nearly identical to that for Case 2 shows that terms (2), (4) and (5) converge to 0, while term (3) converges to 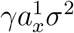.

#### Case 4

0 = *l*_2_ = *l*_1_. The same argument as for term (4) in Case 2 shows that terms (3), (4) and (5) all converge to 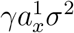. Term (2) is an average of *ε*[*i*]^3^, which is mean zero; consequently, it converges to 0. This completes the proof.

### D Proof of Proposition 5.2

In the limit,

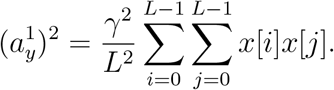

Similarly,

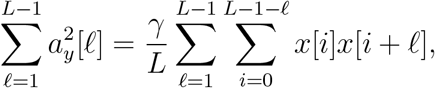

and 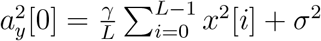. The proof is concluded by noting that 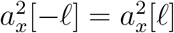.

### E Proof of Proposition 5.3

We prove that both *σ* and *γ* are identifiable from the observed first three moments of *y*. For convenience, we work with *β* = *γ/L* rather than *γ* itself. To this end, we construct two quadratic equations satisfied by *β* and whose coefficients can be computed from observable quantities (in the limit). Then, we show that these equations are independent, and hence that *β* is uniquely defined. Given *β*, we can estimate
*σ* using Proposition 5.2.

Throughout the proof, it is important to distinguish between observed and unobserved values. We denote the observed values by *E_i_* or 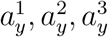. We use *F*_*i*_ to denote functions of the signal’s autocorrelations (which are not directly observable).

In the limit *N* → ∞, almost surely, 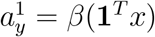 and 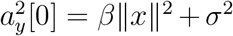, where 1 ∈ ℝ^*L*^ is the vector of all-ones. (In this whole section, for clarity, we now omit to specify that identities hold almost surely in the limit.) Consider the product:

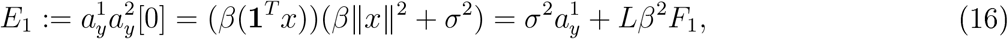

where 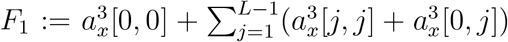. The terms of *F*_1_ can also be estimated from 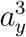, while taking the scaling and bias terms into account. This yields another observable:

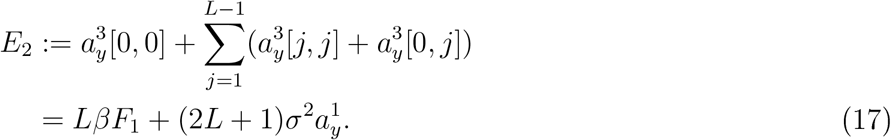

Therefore, from (16) and (17) we get:

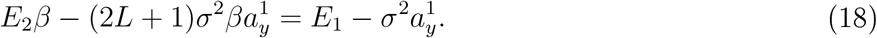

Let 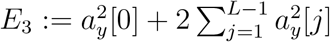; recall from Proposition 5.2:

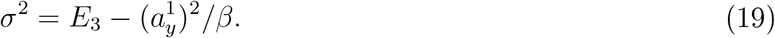

Plugging into (18) and rearranging, we get a first quadratic equation in *β*,

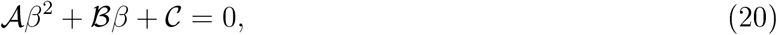

where

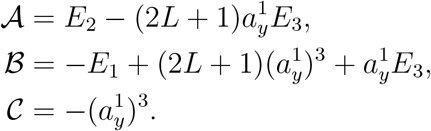

Importantly, these coefficients are observable quantities. As we assume throughout this proof that *x* has nonzero mean, 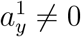 and we conclude that this equation is non-trivial.

Next, we derive the second quadratic equation for *β*. We notice that

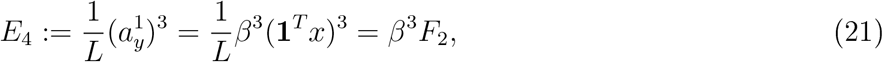

where 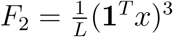, and we can work out that:

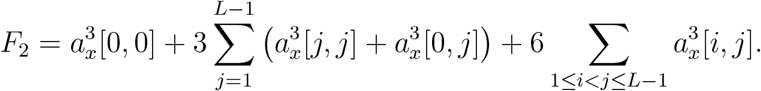

Once again, *F*_2_ can be estimated from 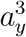, taking bias and scaling into account:

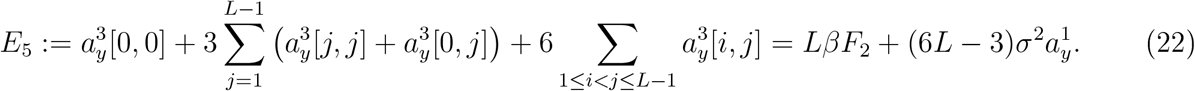

Consider the following ratio:

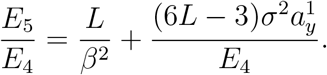

From the latter, we deduce:

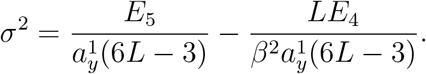

Using (19) and rearranging, we get the second quadratic:

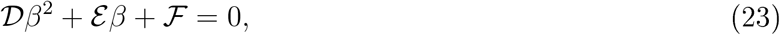

where

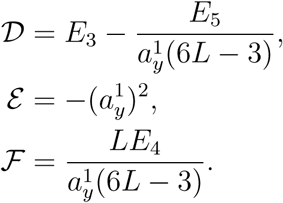

It is also non-trivial since *E*_4_ ≠ 0.

To complete the proof, we need to show that the two quadratic equations (20) and (23) are independent. To this end, it is enough to show that the ratios between coefficients differ. From (20) and (16), we have:

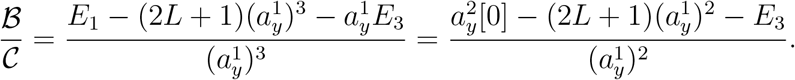

In addition, using (21),

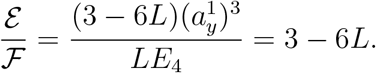

For contradiction, suppose that the quadratics are dependent. Then, 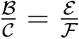, that is,

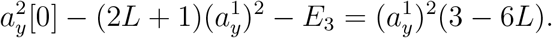

Rewriting the identity in terms of *x* and dividing by *β* we get:

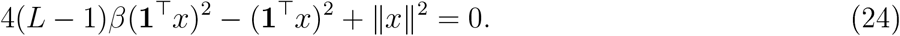

For generic *x*, this polynomial equation is not satisfied so that the quadratic equations are independent. Furthermore, from the inequality *L* ‖*x*‖^2^ ≥ (**1**^⊤^*x*)^2^ it follows immediately that the equations must be independent so long as

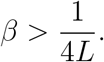

### F Autocorrelations for the cryo-EM problem

#### F.1 Model and autocorrelation functions

Let *ϕ* be the Coulomb potential representing the molecule we aim to recover. We assume that molecule is real-valued and smooth. In spherical coordinates, its 3-D Fourier transform 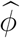 admits a finite expansion of the form

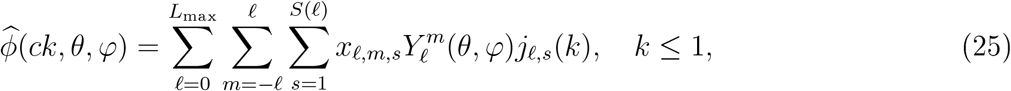

where *c* is the bandlimit, {*S*(*l*)} are determined using the Nyquist criterion as described in [7], *j*_*l,s*_ is the normalized spherical Bessel functions given by

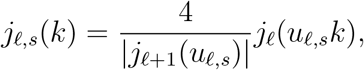

*j*_*l*_ is the spherical Bessel function of order *l* and *u*_*l,s*_ is the *s*th positive zero of *j*_*l*_. We use the complex spherical harmonics 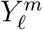 defined by

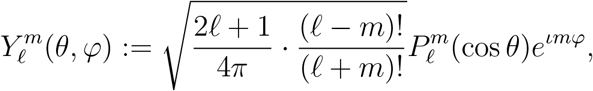

where 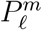 are the associated Legendre polynomials with the Condon-Shortley phase. Sampling at the Nyquist rate dictates *c* = 1/2 [25]. Because *ϕ* is real-valued, 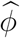 is conjugate-symmetric and thus the expansion coefficients satisfy 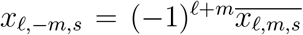. Therefore, we only need to recover coefficients *x*_*l,m,s*_ with *m* ≥ 0.

Let *I*_*ω*_ denote the tomographic projection obtained from viewing direction *ω* ∈ *SO*(3). By the Fourier projection-slice theorem, its 2-D Fourier transform is given by [31]:

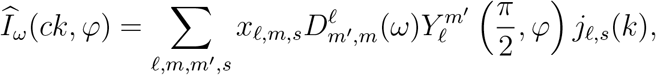

where 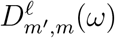 is a Vigner-D matrix. This implies that the projections are also *c*-bandlimited.

Let 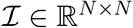 denote a micrograph. We assume it consists of shifted copies of projections contaminated by additive white Gaussian noise:

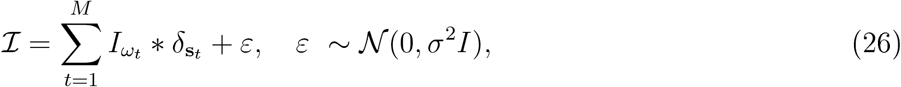

where the viewing directions *ω*_*t*_ are assumed to be drawn from the uniform distribution over SO(3) and s_*t*_ denotes the location of the center of the *t*th projection in the micrograph. We assume the projection is discretized on a Cartesian grid of size *P* × *P* and we impose a separation condition so that any two projections are separated by at least 2*P* − 1 pixels between their upper left corners in each direction, similarly to (2). Note that (1) can be also written as a sum of *δ* functions as in (26).

Define the *p*th autocorrelation of 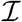 as

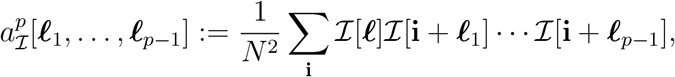

where the summation is for **i** ranging over the *N*^2^ pixels of the micrograph. Let 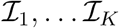 denote a set of *K* micrographs. Under the specified conditions, we show in the next section that the first three autocorrelations of the micrographs are related to those of the projections by

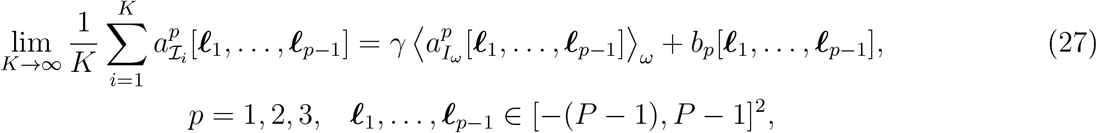

where 〈·〉_*ω*_ denotes averaging over all possible viewing directions *ω* and *b*_*p*_ is a bias term. Specifically, *b*_1_ = 0 and therefore the mean is unbiased. The bias term of the second-order autocorrelation *b*_2_ depends only on *σ*^2^, the variance of the noise. Hence, if the noise level can be accurately estimated from the micrographs, this bias can be removed. Finally, the bias term of the third-order autocorrelation *b*_3_ depends on the mean of the micrograph and *σ*^2^. Therefore, given sufficiently many projections, we can accurately estimate the quantities 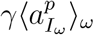 directly from the micrographs. These quantities are functions of the unknown coefficients *x*_*l,m,s*_ and we could proceed to invert their relation, as we did in the toy examples.

In practice, we want to leverage one more feature of the 3-D reconstruction problem. Since all in-plane rotations of the micrographs are equally likely observations, it is desirable in (27) to average over all in-plane rotations as well. This can be done efficiently using Prolate Spheroidal Vave Functions (PSVFs). We use autocorrelations up to and including the third order. Indeed, second-order autocorrelations are not enough, as was observed already in [21] for a simpler problem where the input is not micrographs but rather picked, perfectly centered particles.

#### F.2 Autocorrelation derivation

In this section we prove relation (27). We note that mathematically taking infinitely many micrographs is equivalent to take one infinitely large micrograph with fixed density *γ*. Hence, we consider the moments of one micrograph 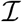 in the limit *N* → ∞ and 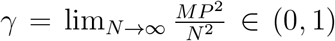. The separation condition
guarantees that if **i** = (*i, j*) is in the support of some projection, then **i** + ***l*** for ***l*** ∈[−(*P* − 1), *P* − 1]^2^ is either in the support of the same projection or outside the support of any projection.

We begin by calculating the relation between the *p*th autocorrelation of the clean micrograph and the averaged autocorrelation of the projections. Let us denote the clean micrograph by 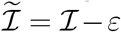, where 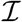 and *ε* are given in (26). Denote by 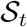 the support of the *t*th particle. Then, we have

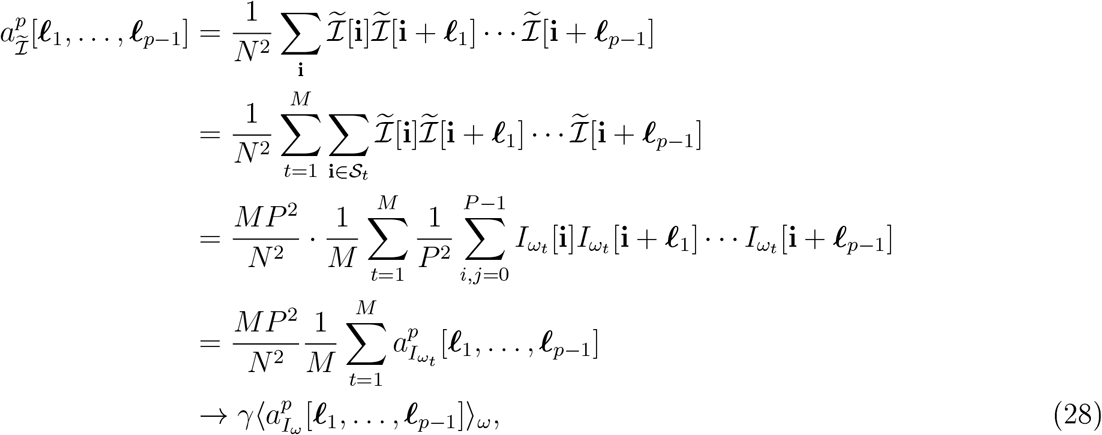

where the average is taken over *ω* with respect to the distribution of viewing directions. Here, we assume it to be uniform.

In the presence of noise, we get additional bias terms denoted by *b*_*p*_ in (27). The mean (*p* = 1) is unbiased since the noise is assumed to have zero mean. For the second-order autocorrelation (*p* = 2), we have

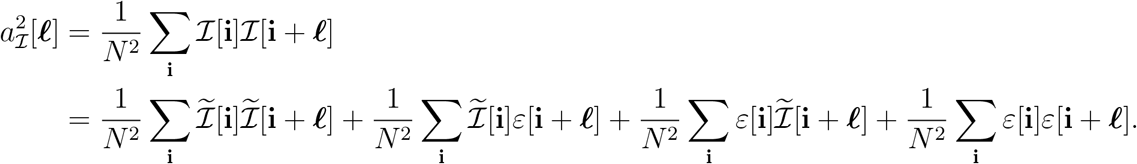

The first term is given by (28) for *p* = 2. The cross terms vanish in the limit. The fourth term is zero unless *l* = 0, in which case it converges to *σ*^2^. Thus, we conclude

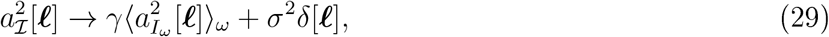

where the bias term *b*_2_[***l***] = *σ*^2^*δ*[***l***] depends only on the variance of the noise *σ*^2^.

For the third moments, we get 8 terms:

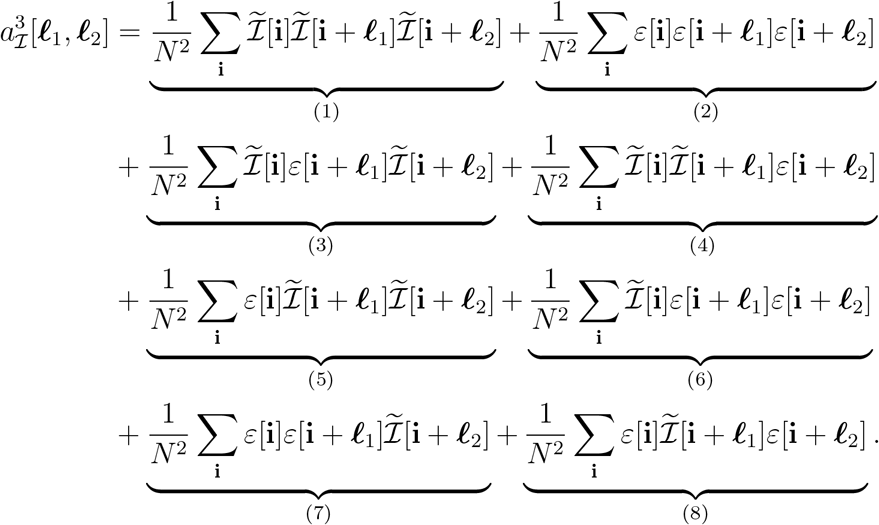

We address these terms one by one:

- Term (1) is treated by (28) for *p* = 3;
- Term (2) is the third-order autocorrelation of pure noise which vanishes in the limit;
- Terms (3)–(5) depend linearly on the noise and hence vanish in the limit;
- For term (6), if ***l***_1_ ≠ ***l***_2_ the term vanishes in the limit. If ***l***_1_ = ***l***_2_ then

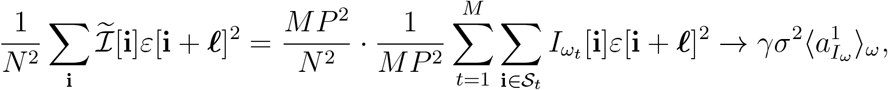

where 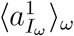 is the mean of the volume.
- Terms (7) and (8) contribute *δ* functions similar to (6). Thus, we conclude that

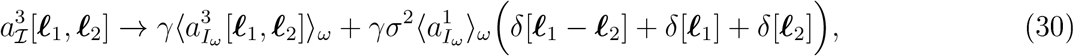

where the second term is the bias *b*_3_[***l***_1_, ***l***_2_]. Note that 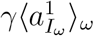 is approximately the mean of the micrograph since 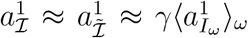. Therefore, we do not need prior knowledge of *γ* to effectively debias the third-order autocorrelation.

#### F.3 Accounting for all in-plane rotations

We represent our autocorrelations using Prolate Spheroidal Vave Functions (PSVFs) {*Ψ*_*k,q*_} where *k* ≥ 0, *q* ≥ 1 are integers [39]. As we demonstrate below, this makes it easier to account for the fact that all in-plane rotations of the micrographs are equally likely observations. This is only a concern for the second and third order autocorrelations. Below, we start with *p* = 2. The PSVFs are eigenfunctions of the truncated Fourier transform and are given in polar coordinates by^5^

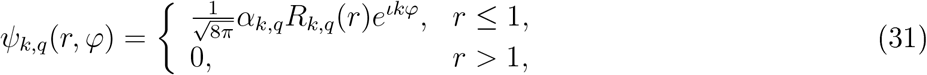

where the range of *k, q* is determined by Eq. (8) in [24], the *R*_*k,q*_ are a family of real, one-dimensional functions and the *α*_*k,q*_ are scaling factors which will be defined in the next section. The PSVFs are orthogonal on the unit disk.

For ***l*** ∈ [−(*P* − 1), *P* − 1]^2^, let us define

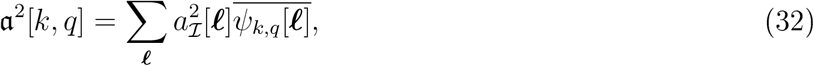

where *Ψ*_*k,q*_[***l***] := *Ψ*_*k,q*_(***l***/(*P* − 1)) is a discretization of the PSVFs. Knowledge of these coefficients is essentially equivalent to knowledge of the second-order correlations owing to the following approximate identity:

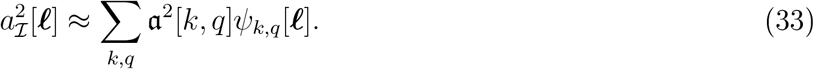

This holds because the continuous PSVFs form an orthogonal basis, and their discretized counterparts are (empirically) almost orthogonal. As a result, for our purposes, the pair of equations above provides a basis expansion for the autocorrelations.

We now proceed to show that the coefficients *a*^2^[*k, q*] can be computed from the micrographs directly.

By definition,

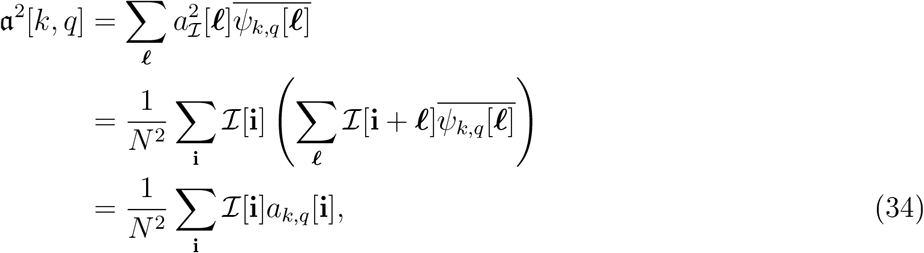

where we defined

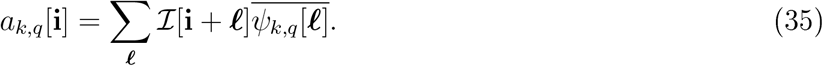

These coefficients can be computed efficiently. Indeed, consider a patch of the micrograph centered around pixel 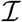 and of size (2*P*− 1) × (2*P* − 1). This is exactly the patch indexed in the sum above. Hence, using the same approximation as we did in (33), a direct expansion of that patch in the discretized PSVFs yields the sought coefficients:

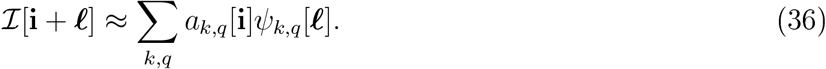

Thus, we proceed as follows: for each position **i** in the micrograph 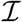, we extract the corresponding patch of size (2*P* − 1) × (2*P* − 1), expand it in the discretized PSVFs as in (36), and collect the *a_k,q_* as per (34) to constitute the second-order autocorrelation of the micrograph.

Crucially, following this formalism, it is now straightforward to account for all in-plane rotations and reflections of the micrograph. Indeed, as can be seen from the definition of the (continuous) PSVFs (31), the effects of rotations and reflections on expansion coefficients of real images are, respectively, phase modulation and conjugation: this is why this the PSVF basis is called *steerable* [24, 45]. By analogy in the discrete case, we have the following approximate expansions for a patch rotated about its center **i** by an angle *α*:

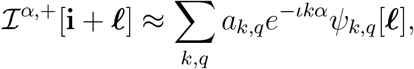

and the reflection followed by a rotation by angle *α*:

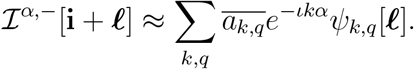

Averaging over all rotations of the patch 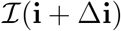 and its reflection we get

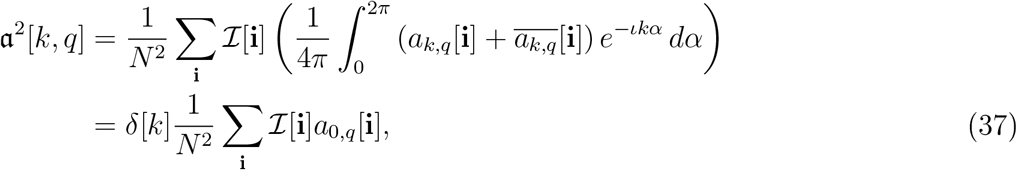

where in the last equality we used that *a*_0,*q*_[**i**] is real since both 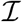 and *ѱ*_0,*q*_ are real valued (more generally, 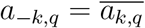). Thus, the second-order autocorrelation, though two-dimensional, effectively only provides radial information.

We now follow a similar approach to estimate the bias term *b*_2_. Introduce the coefficients *b*_2_ as:

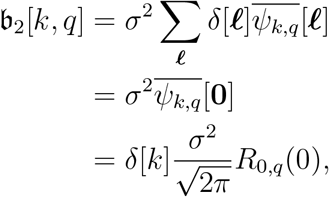

where we used the fact that the functions *R*_*k,q*_ are zero at the origin for *k* ≠ 0. With this definition, we have the usual approximation:

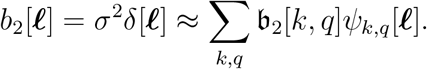

We now turn out attention to the third order autocorrelation. Following the same lines, we define the coefficients:

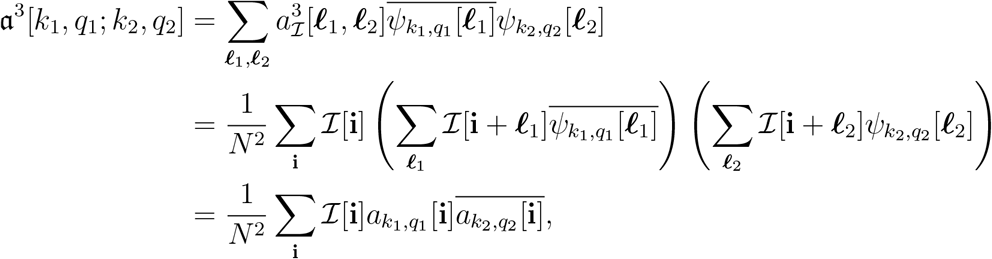

where the patch expansion coefficients *a*_*k,q*_ are as defined in (35). The coefficients *a*^3^[*k*_1_, *q*_1_; *k*_2_, *q*_2_] are related to the third-order autocorrelation via the approximate identity:

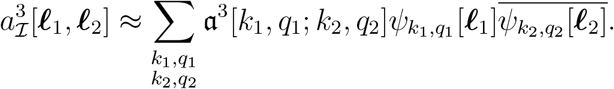

Averaging over all rotations of 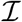 and its reflection, we obtain

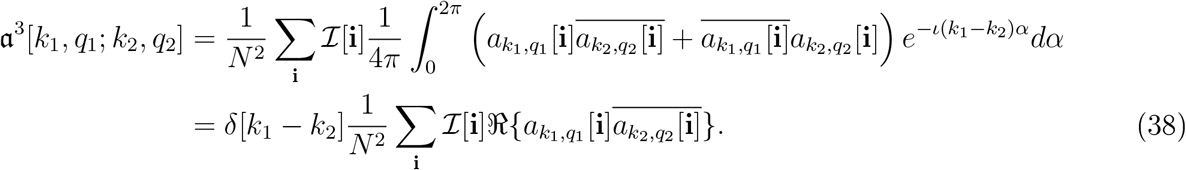

Thus, similarly to the second-order autocorrelation, averaging over all in-plane rotations reveals that the 4-D third-order autocorrelation truly only carries information along three dimensions.

Finally, we treat the bias terms:

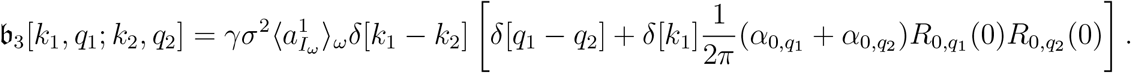

Thus,

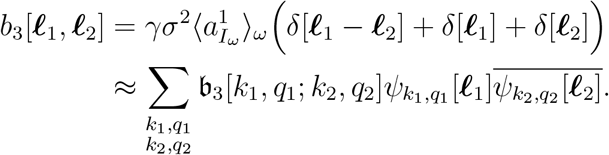

#### F.4 Connection to volume

Until now, we have established simple relations between the autocorrelations of the micrographs and the autocorrelations of the volume. Now, we complete the picture by deriving the connection with the volume itself.

Using the 2-D PSWFs, we can express each projection as

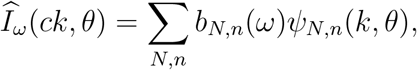

where

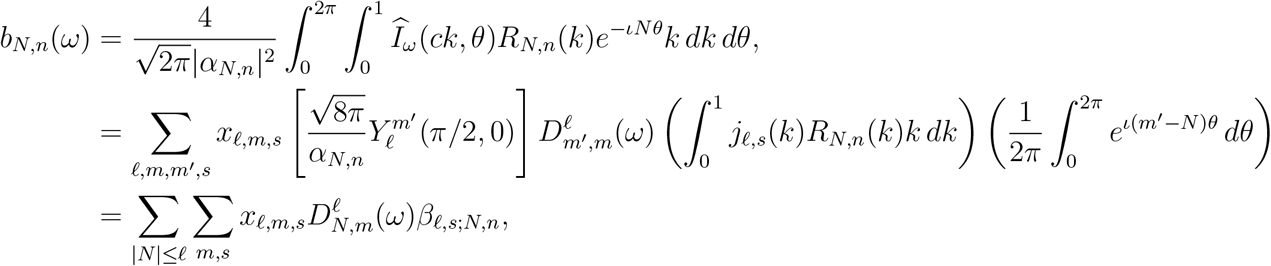

where the coefficients

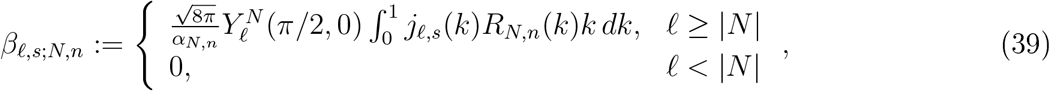

can be precomputed. Since the PSWFs are eigenfunctions of the truncated Fourier transform [24] and hence satisfy

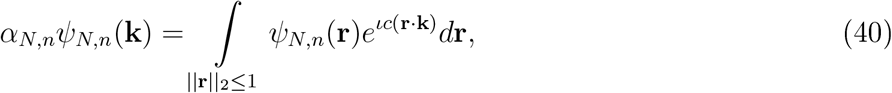

we can now express the projection in real space as

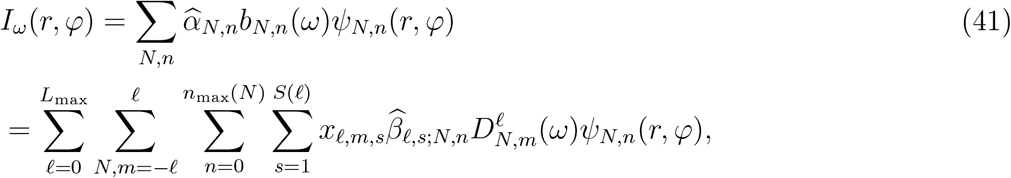

where *n*_*max*_(*N*) is chosen according to Eq. (8) in [24], *α*_*N,n*_ is the eigenvalue corresponding to the (*N, n*)th PSWF, 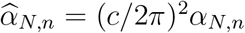, and 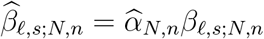.

##### Third-order autocorrelation

In (30) and (38) we have shown how the third-order autocorrelations of the micrographs and the volume are related, and how we can present them in PSVFs. Now, we relate these expressions to the expansion coefficients of the volume itself.

The third-order autocorrelation of the volume can be expressed in terms of (41):

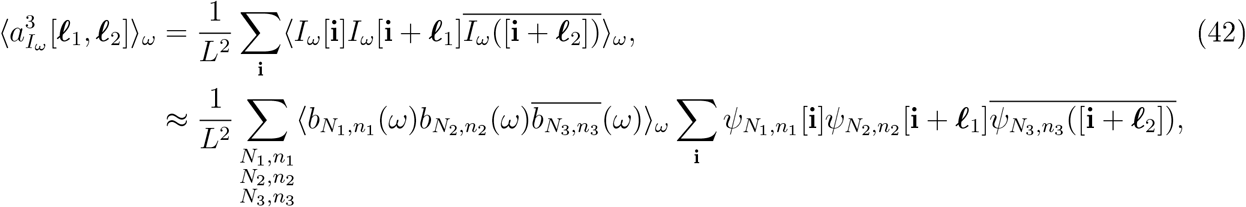

where the approximation is due to discretization. Now,

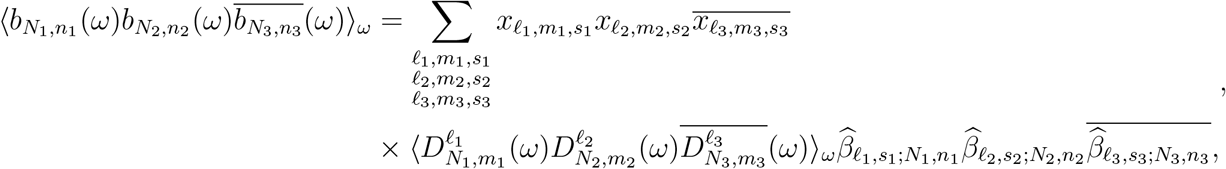

where the latter coefficients are given explicitly in (39). Using standard properties of D-Wigner functions, we obtain

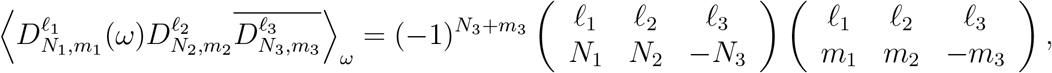

where 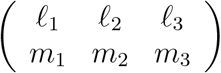 are called Wigner 3-j symbols. Notably, these terms are zero unless *m*_1_ + *m*_2_ + *m*_3_ = 0 and |*l*_1_ − *l*_2_| ≤ *l*_3_ ≤ *l*_1_ + *l*_2_. Thus, we conclude that

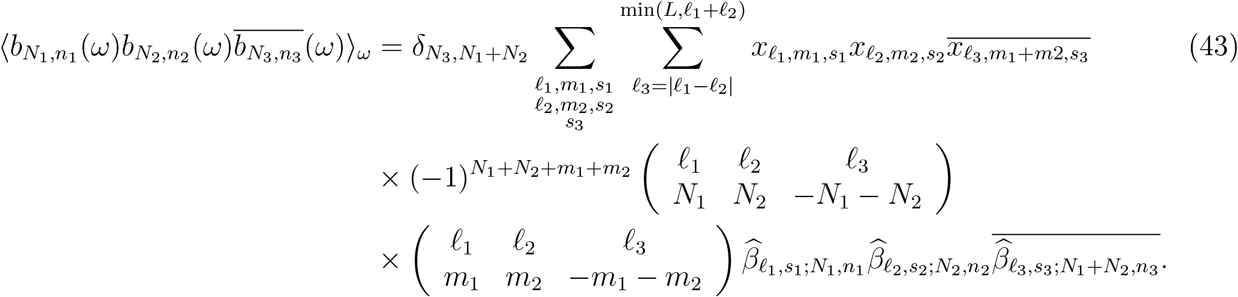

Combining (43) with (42) provides the explicit relation between the third-order autocorrelation and the volume.

Recall that we obtain the autocorrelations of the volume in PSWFs coefficients; see (38). Hence, to conclude the derivation we expand

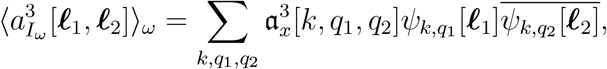

where we only include the block-diagonal terms in the expansion; the rest are equal to zero. Let

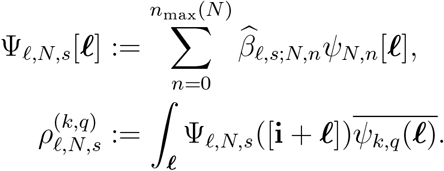

Then, the final formula reads

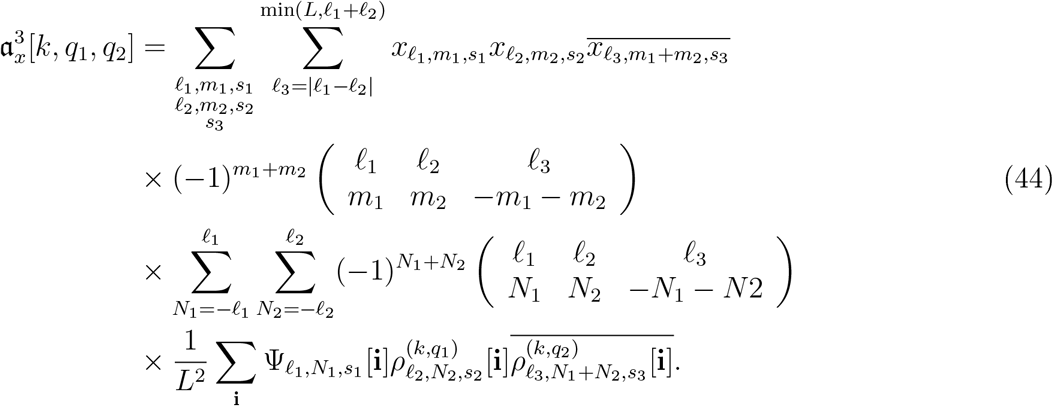

In practice, the last two lines of the above expression for 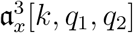 is precomputed, and both the integration over **i** and over ***ℓ*** is performed on the grid of the images in the dataset, to match the integration performed on the actual images.

##### Second-order autocorrelation

The second-order autocorrelation is easier to derive directly in Fourier space, to avoid integration of shifted PSVFs against centered ones. The relation between the second-order autocorrelation of the micrographs and the volume is given in (29) and (37). The connection with the expansion coefficients of the volume can be derived in Fourier space directly from Kam’s original formula [21] by setting **k**_1_ = **k**_2_ to obtain

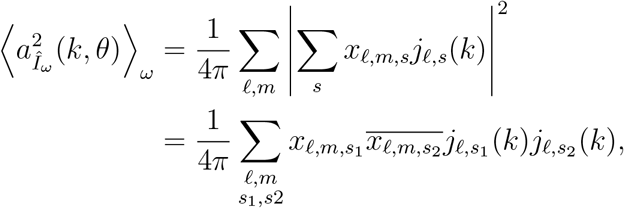

where we used the fact that the normalized spherical Bessel functions *j*_*l,s*_ are real.

As before, we want to derive the relation with respect to the PSVF coefficients of the autocorrelation.
Hence, we expand the above in 2-D PSWFs by

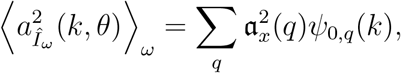

and conclude that

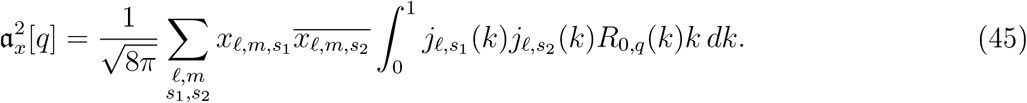

The integral on *k* is precomputed.

##### The mean

Since *j*_*l,s*_(0) = 0 unless *l* = 0, and since 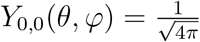, we conclude from (25) that

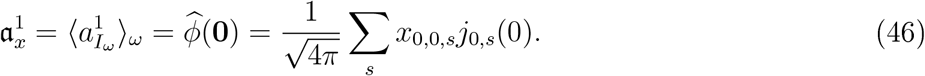

## Footnotes

1 Autocorrelation in *d* dimensions is defined similarly to (4), where *i,l*_1_, …, *l*_*q* − 1_ take on values in ℤ^*d*^ and the summation in *i* is over all of ℤ^*d*^.

2 https://www.rcsb.org

3 http://www.ebi.ac.uk/pdbe/emdb

4 http://spr.math.princeton.edu

5 A different normalization is used in [24].

## References

[1] Emmanuel Abbe, Tamir Bendory, William Leeb, João Pereira, Nir Sharon, and Amit Singer, Multireference alignment is easier with an aperiodic translation distribution, arXiv preprint arXiv.l710.02793, 2017.

[2] Karim Abed-Meraim, Wanzhi Qiu, and Yingbo Hua, Blind system identification. Proceedings of the IEEE., 85(8): 1310–1322, 1997.

[3] GE Ayers and J Christopher Dainty. Iterative blind deconvolution method and its applications. Optics letters, 13(7):547–549, 1988.

[4] Afonso S Bandeira, Ben Blum-Smith, Amelia Perry, Jonathan Weed, and Alexander S Wein. Estimation under group actions: recovering orbits from invariants. arXiv preprint arXiv:1712.10163, 2017.

[5] Tamir Bendory, Nicolas Boumal, Chao Ma, Zhizhen Zhao, and Amit Singer. Bispectrum inversion with application to multireference alignment. IEEE Transactions on Signal Processing, 66(4): 1037–1050, 2017.

[6] Albert Benveniste, Maurice Goursat, and Gabriel Euget. Eobust identification of a nonminimum phase system: Blind adjustment of a linear equalizer in data communications. IEEE Transactions on Automatic Control, 25(3):385–399, 1980.

[7] Tejal Bhamre, Teng Zhang, and Amit Singer. Anisotropic twicing for single particle reconstruction using autocorrelation analysis. arXiv preprint arXiv:1704-07969, 2017.

[8] N. Boumal, B. Mishra, P.-A. Absil, and E. Sepulchres. Manopt, a Matlab toolbox for optimization on manifolds. Journal of Machine Learning Research, 15:1455–1459, 2014.

[9] Nicolas Boumal, Tamir Bendory, Roy R Lederman, and Amit Singer, Heterogeneous multireferenee alignment: A single pass approach. In Information Sciences and Systems (CISS), 2018 52nd Annual Conference on, pages 1–6, IEEE, 2018.

[10] Olivier Cappé, Arnaud Doucet, Marc Lavielle, and Eric Moulines, Simulation-based methods for blind maximum-likelihood filter identification. Signal processing, 73(1-2):3—25, 1999.

[11] David Donoho, Jiashun Jin, et al. Higher criticism for detecting sparse heterogeneous mixtures. The Annals of Statistics, 32(3):962–994, 2004.

[12] Veit Elser, Matrix product constraints by projection methods. Journal of Global Optimization, 68(2):329–355, 2017.

[13] Joachim Frank, Single-particle reconstruction of biological moleeules-storv in a sample (Nobel lecture), Angewandte Chemie International Edition, 2018.

[14] Georgios B Giannakis and Jerry M Mendel, Identification of nonminimum phase systems using higher order statistics, IEEE Transactions on Acoustics, Speech, and Signal Processing, 37(3):360–377, 1989.

[15] Robert M Glaeser. Electron crystallography: present excitement, a nod to the past, anticipating the future. Journal of structural biology, 128(1):3–14, 1999.

[16] Sandeep Gogineni, Pawan Setlur, Muralidhar Rangaswamv, and Raj Rao Nadakuditi. Passive radar detection with noisy reference channel using principal subspace similarity. IEEE Transactions on Aerospace and Electronic Systems, 54(1):18—36, 2018.

[17] Mhmh Hayes. The reconstruction of a multidimensional sequence from the phase or magnitude of its Fourier transform. IEEE Transactions on Acoustics, Speech, and Signal Processing, 30(2):140–154, 1982.

[18] Richard Henderson. The potential and limitations of neutrons, electrons and X-rays for atomic resolution microscopy of unstained biological molecules. Quarterly reviews of biophysics, 28(2):171–193, 1995.

[19] Richard Henderson. Avoiding the pitfalls of single particle ervo-eleetron microscopy: Einstein from noise. Proceedings of the National Academy of Sciences, 110(45):18037–18041, 2013.

[20] Stuart M Jefferies and Julian C Christou. Restoration of astronomical images by iterative blind deconvolution. The Astrophysical Journal, 415:862, 1993.

[21] Zvi Kam. The reconstruction of structure from electron micrographs of randomly oriented particles. Journal of Theoretical Biology, 82(1): 15–39, 1980.

[22] Maryam Khoshouei, Mazdak Radjainia, Wolfgang Baumeister, and Radostin Danev, Crvo-EM structure of haemoglobin at 3,2 A determined with the Volta phase plate. Nature communications, 8:16099, 2017.

[23] John Kormvlo and J Mendel. Identifiabilitv of nonminimum phase linear stochastic systems. IEEE transactions on automatic control, 28(12):1081–1090, 1983.

[24] Boris Landa and Yoel Shkolniskv, Steerable principal components for spaee-frequenev localized images. SIAM journal on imaging sciences, 10(2):508–534, 2017.

[25] Eitan Levin, Tamir Bendory, Nicolas Boumal, Joe Kileel, and Amit Singer, 3D ab initio modeling in ervo-EM by autocorrelation analysis. In Biomedical Imaging (ISBI 2018), 2018 IEEE 15th International Symposium on, pages 1569–1573, IEEE, 2018.

[26] Michael S Lewieki, A review of methods for spike sorting: the detection and classification of neural action potentials. Network: Computation in Neural Systems, 9(4):E53–E78, 1998.

[27] Yi-Lvnn Liang, Maryam Khoshouei, Mazdak Eadjainia, Yan Zhang, Alisa Glukhova, Jeffrey Tar-rasch, David M Thai, Sebastian GB Furness, George Christopoulos, Thomas Coudrat, et al, Phase-plate ervo-EM structure of a class B GPCE-G-protein complex. Nature, 546(7656): 118, 2017.

[28] Ks Lii, M Rosenblatt, et al. Deconvolution and estimation of transfer function phase and coefficients for nongaussian linear processes. The annals of statistics, 10(4); 1195—1208, 1982.

[29] Yuxi Liu, Shane Gonen, Tamir Gonen, and Todd O Yeates, Near-atomic ervo-EM imaging of a small protein displayed on a designed scaffolding system. Proceedings of the National Academy of Sciences, 115(13):3362–3367, 2018.

[30] Lennart Ljung, System identification. In Signal analysis and prediction, pages 163–173, Springer, 1998.

[31] Frank Natterer. The mathematics of computerized tomography, volume 32. SIAM, 1986.

[32] Eric F Pettersen, Thomas D Goddard, Conrad C Huang, Gregory S Couch, Daniel M Greenblatt, Elaine C Meng, and Thomas E Ferrin. UCSF chimera—a visualization system for exploratory research and analysis. Journal of computational chemistry, 25(13) :1605–1612, 2004.

[33] Lawrence E Eabiner. A tutorial on hidden Markov models and selected applications in speech recognition. Proceedings of the IEEE, 77(2):257–286, 1989.

[34] Giovanna Scapin, Clinton S Potter, and Bridget Carragher. Crvo-EM for small molecules discovery, design, understanding, and application. Cell Chemical Biology, 2018.

[35] Sjors HW Scheres. RELION: implementation of a bavesian approach to ervo-EM structure determination. Journal of structural biology, 180(3) :519–530, 2012.

[36] Ofir Shalvi and Ehud Weinstein. New criteria for blind deconvolution of nonminimum phase systems (channels). IEEE Transactions on information theory, 36(2):312–321, 1990.

[37] Maxim Shatskv, Eichard J Hall, Steven E Brenner, and Eobert M Glaeser. A method for the alignment of heterogeneous macromolecules from electron mieroseopv. Journal of structural bioloqy, 166(1) :67–78, 2009.

[38] FJ Sigworth. A maximum-likelihood approach to single-particle image refinement. Journal of structural biology, 122(3):328–339, 1998.

[39] David Slepian. Prolate spheroidal wave functions, Fourier analysis and uncertainty—iv: Extensions to many dimensions; generalized prolate spheroidal functions. The Bell System Technical Journal, 43(6):3009–3057, Nov 1964.

[40] Gilbert W Stewart. The efficient generation of random orthogonal matrices with an application to condition estimators. SIAM Journal on Numerical Analysis, 17(3):403–409, 1980.

[41] Jitendra Tugnait, Identification of nonminimum phase linear stochastic systems. In The 23rd IEEE Conference on Decision and Control, number 23, pages 342–347, 1984.

[42] Marin van Heel, Finding trimeric HIV-1 envelope glycoproteins in random noise. Proceedings of the National Academy of Sciences, 110(45):E4175–E4177, 2013.

[43] Marin van Heel, Michael Schatz, and Elena Orlova, Correlation functions revisited. Ultramicroscopy, 46(1-4) :307–316, 1992.

[44] Yuqian Zhang, Han-Wen Kuo, and John Wright, Structured local optima in sparse blind deconvolution, arXiv preprint arXiv:1806.00338, 2018.

[45] Zhizhen Zhao, Yoel Shkolnisky, and Amit Singer, Fast steerable principal component analysis, IEEE transactions on computational imaging, 2(1): 1—12, 2016.

